# Mono-phosphorylation at Ser4 of Barrier-to-autointegration Factor (Banf1) significantly reduces its DNA binding capability by inducing critical changes in its local conformation and DNA binding surface

**DOI:** 10.1101/2023.05.21.541616

**Authors:** Ming Tang, Amila Suraweera, Xuqiang Nie, Zilin Li, James W. Wells, Kenneth J O’Byrne, Robert J Woods, Emma Bolderson, Derek J Richard

## Abstract

Barrier-to-Autointegration Factor (Banf1) is a small DNA-bridging protein. The binding status of Banf1 to DNA is regulated by its N-terminal phosphorylation and dephosphorylation, which plays a critical role in cell proliferation. Banf1 can be phosphorylated at Ser4 into mono-phosphorylated Banf1, which is further phosphorylated at Thr3 to form di-phosphorylated Banf1. It was observed decades ago that mono-phosphorylated Banf1 cannot bind to DNA. However, the underlying molecular- and atomic-level mechanisms remain unclear. A clear understanding of these mechanisms will aid in interfering with the cell proliferation process for better global health. Herein, we explored the detailed atomic bases of unphosphorylated Banf1-DNA binding and how mono- and di-phosphorylation of Banf1 impair these atomic bases to eliminate its DNA-binding capability, followed by exploring the DNA-binding capability of mono- and di-phosphorylation Banf1, using comprehensive and systematic molecular modelling and molecular dynamics simulations. This work presented in detail the residue-level binding energies, hydrogen bonds and water bridges between Banf1 and DNA, some of which have not been reported. Moreover, we revealed that mono-phosphorylation of Banf1 causes its N-terminal secondary structure changes, which in turn induce significant changes in Banf1’s DNA binding surface, thus eliminating its DNA-binding capability. At the atomic level, we also uncovered the alterations in interactions due to the induction of mono-phosphorylation that result in the N-terminal secondary structure changes of Banf1. Additionally, our modelling showed that phosphorylated Banf1 with their dominant N-terminal secondary structures bind to DNA with a significantly lower affinity and the docked binding pose are not stable in MD simulations. These findings help future studies in predicting effect of mutations in Banf1 on its DNA-binding capability and open a novel avenue for the development of therapeutics such as cancer drugs, targeting cell proliferation by inducing conformational changes in Banf1’s N-terminal domain.

## Introduction

Barrier-to-Autointegration Factor 1 (Banf1) is a small protein with a monomer mass of 10 kDa, consisting of 89 residues [1]. Banf1 is highly conserved in metazoans [2]. Through interacting with DNA, histones and many other nuclear proteins, Banf1 plays an indispensable role in cell proliferation [3]. Banf1 is also involved in numerous other cellular processes [4–8]. We have recently reported that Banf1 binds to and regulates the DNA repair activity of Poly [ADP-ribose] Polymerase 1 (PARP1) and DNA-dependent Kinase (DNA-PK) following DNA damage [7, 8].

Banf1 monomers form stable homodimers and ultimately oligomers, enabling them to bind strongly to double-stranded DNA (dsDNA, hereafter denoted as DNA) [4]. This further permits Banf1 to bridge and condense intra- and/or inter-molecular DNA helices [9] within foreign and genomic DNA [4, 10], which is critical to most of Banf1’s fundamental cellular functional roles in protecting genome integrity and ensuring successful completion of mitosis [3]. Unphosphorylated Banf1 binds to DNA as a dimer, with each of the two protein monomers bound to one DNA molecule (Figure 1A). As illustrated in Figure 1, X-ray crystallography study revealed that DNA binding by Banf1 is achieved by: i) a water bridge between the side chain of Ser4 and the backbone phosphate of DA6; ii) Van der Waals interaction between the sidechain of Val29 and the ribose of DC5; iii) seven hydrogen bonds between DNA phosphate backbone and the backbone of Banf1 residues, including Gly25, Gly27, Val29 and Leu30 at helix-hairpin-helix (HhH) motif (in pink), Ala71 at pseudo HhH motif (in blue), and Gln5 and Lys6 at the N-terminal α1 helix (in yellow); and iv) two hydrogen bonds between DNA phosphate backbone and the sidechain of Banf1 residues, including guanidine group of Arg75 and ε-amino group of Lys6 [11]. Notably, in the co-crystalised Banf1_Thr2_-DNA (PDB ID: 2BZF), there are other Banf1 hydrophobic residues (e.g., Ala24 and Ile26) and polar residues (e.g., Asn70, Gln73, and Lys72) located within 5 Å from DNA, which may form hydrophobic interactions with the ribose of DNA nucleotides and hydrogen bonding interactions and/or water bridges with DNA nucleotides, respectively. Therefore, it is important to explore all residues in Banf1 that contribute to its DNA binding and to quantify the energy contribution of both their backbone and sidechain, which provides precious insights into whether the numerous mutations found in Banf1 may affect its DNA binding capability and hence cellular function [12, 13].

**Figure 1.**
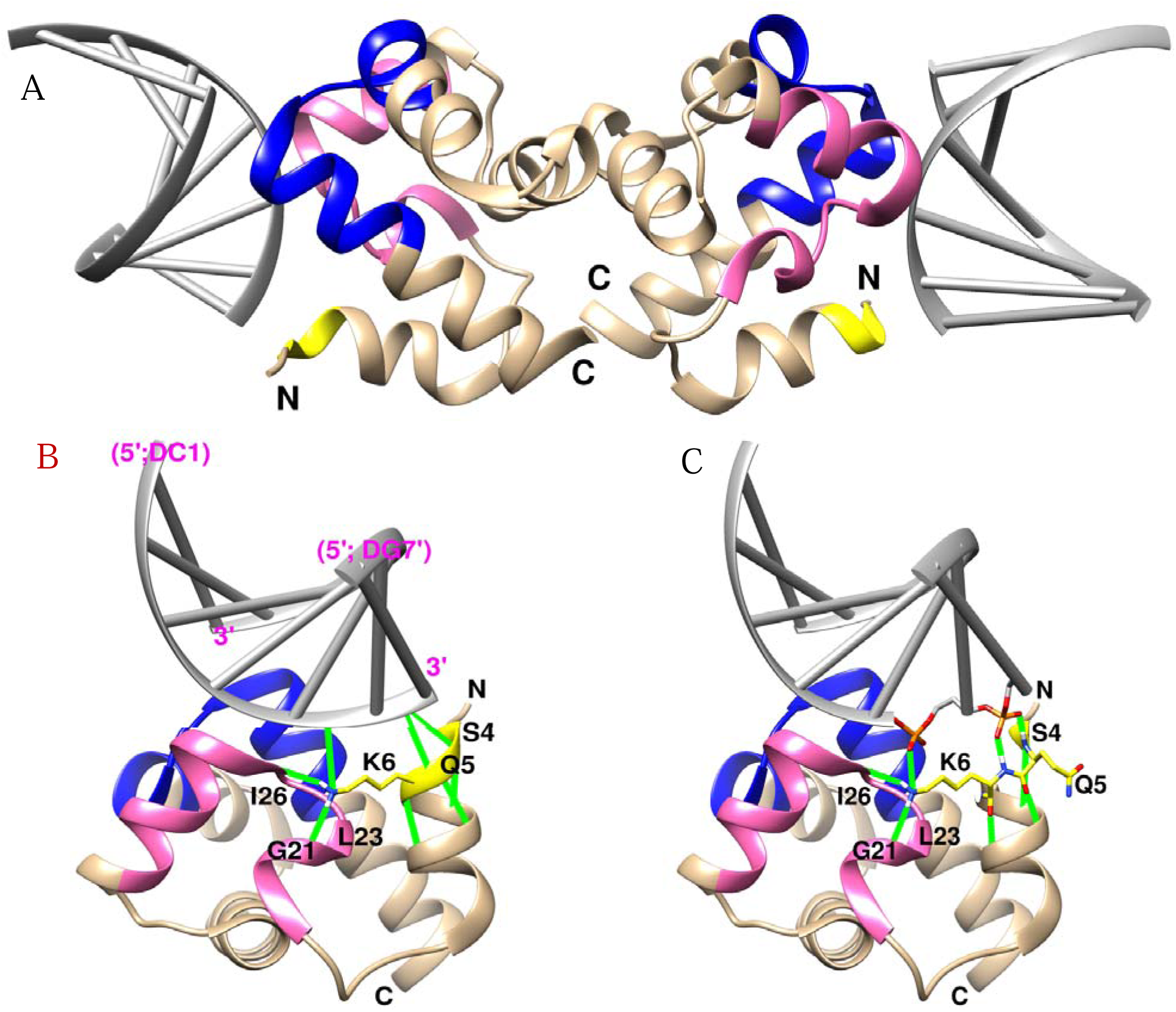
Molecular structure of Banf1 in complex with DNA double helix (PDB ID: 2BZF) coloured in dark gray. A. Pure ribbon representation of Banf1 dimer complexed with two DNA molecules. B. Pure ribbon representation of Banf1 monomer complexed with a DNA double helix. C. Ribbon representation of the complex in B, with Banf1’s N-terminal residues 5 and 6 and the DNA phosphate backbones they interact with presented with sticks. Banf1 monomer interacts with DNA via the helix-hairpin-helix motif (hot pink), the pseudo helix-hairpin-helix motif (blue), and the N-terminal region (yellow), which are at the opposite side of the dimerization surface. The backbone amide nitrogens of Gln5 and Lys6 hydrogen bond to DNA phosphate oxygens. The ε-amino group of Lys6 is restricted by the carbonyl groups of Gly21, Leu23 and Ile26 and forms a salt bridge with DNA phosphate oxygen. Hydrogen bonds and salt bridges are displayed as green lines. Figure is generated in Chimera 1.15. Nucleotides are numbered 1 to 7 in the 5’-to-3’ direction of the DNA strand that interacts with Banf1 N-terminal domain and 1’ to 7’ in the 3’-to-5’ direction of the other strand.

Unlike unphosphorylated Banf1 that strongly binds to DNA [11], both mono-phosphorylated Banf1 [14–16] and di-phosphorylated Banf1 [14, 15] were found unable to bind DNA or binds DNA with a significantly lower affinity. Banf1 is initially phosphorylated at Ser4 forming mono-phosphorylated Banf1, which can be further phosphorylated at Thr3 (pThr3), resulting in di-phosphorylated Banf1 [17]. The ability of Banf1 to bind to DNA is severely inhibited by phosphorylation of its N-terminal residue(s) [17, 18]. Specifically, the binding affinity of unphosphorylated Banf1 for DNA is 2.5 ± 1 nM, which is thousands of times higher than the binding affinity (11 ± 2 μM) of di-phosphorylated Banf1 for DNA [17]. Interestingly, the N-terminal secondary structure reported for the di-phosphorylated Banf1 dimer alone and for the di-phosphorylated Banf1 dimer co-crystalised with the LEM domain of emerin are relatively different; the N-terminal α1 helix of di-phosphorylated Banf1 dimer co-crystalised with emerin (PDB ID: 7NDY) is elongated to residues 2-11, whereas the N-terminal α1 helix in di-phosphorylated Banf1 dimer alone is shortened to residues 7-11 (Figure 5A in Ref. [17]). Therefore, we speculate that the secondary structure of the N-terminal region of di-phosphorylated Banf1 may be relatively dynamic. Notably, it has been reported that phosphorylation at Ser4 of Banf1 alone is sufficient to eliminate its DNA-binding capability [17, 18].

The detailed atomic bases of unphosphorylated Banf1-DNA binding are not fully understood. Moreover, the dynamics of the secondary structure of the N-terminal domain of di-phosphorylated Banf1 are not fully understood. Additionally, the molecular- and atomic-level mechanisms that define how mono-phosphorylation of Banf1 impairs its DNA-binding capability remain unclear, although the phenomenon has been observed and reported decades ago. To fill these research gaps, we used molecular docking and molecular dynamics simulations i) to uncover the molecular bases underlying the Banf1-DNA binding; ii) to explore the impact of mono- and di-phosphorylation on the N-terminal secondary structure of Banf1 and its DNA binding surface; and iii) to explore the DNA-binding capability of mono- and di-phosphorylation Banf1. The computational modelling results revealed that mono-phosphorylation of Banf1 inhibits its binding to DNA, by inducing steric clashes and unfavourable interactions originating from distinct changes in local N-terminal conformational and DNA-binding surface of Banf1. Similar molecular modelling techniques have been applied to successfully unravel the molecular- and/or atomic-level mechanisms of biological systems that were challenging for then-current experimental techniques [19–24], including designing engineered transmembrane proteins recently reported by us [25], binding of transcription factors to DNA [26], complicated dynamics of enzymes [27, 28], importance of WPD-Loop sequence for activity and structure in protein tyrosine phosphatases [29], the surface adhesion mechanisms of COVID-19 [30], the nanomechanical features of coronavirus spike proteins [31, 32], and the conformational dynamics of proteins related to diseases [33–39].

### Computational models and methodology

#### Model building

Banf1 exists in cells in two forms, i.e., Banf1 with a methionine residue at the initial position in the sequence (Banf1_Met1_) and without the methionine, in which case the sequence begins with threonine (Banf1_Thr2_). Herein, we performed systematic and comprehensive MD simulations of i) Banf1-DNA complexes, including Banf1_Met1_-DNA, Banf1_Thr2_-DNA, mono-phosphorylated Banf1_Met1_-DNA (pSer4 Banf1_Met1_-DNA), di-phosphorylated Banf1_Met1_-DNA (pBanf1_Met1_-DNA), mono-phosphorylated Banf1_Thr2_-DNA (pSer4 Banf1_Thr2_-DNA) and di-phosphorylated Banf1_Thr2_-DNA (pBanf1_Thr2_-DNA); ii) Banf1 dimers, including unphosphorylated Banf1_Met1_ (WT Banf1_Met1_), mono-phosphorylated Banf1_Met1_ (pSer4 Banf1_Met1_), di-phosphorylated Banf1_Met1_ (pBanf1_Met1_), unphosphorylated Banf1_Thr2_ (WT Banf1_Thr2_), mono-phosphorylated Banf1_Thr2_ (pSer4 Banf1_Thr2_) and di-phosphorylated Banf1_Thr2_ (pBanf1_Thr2_); and iii) Banf1 monomers of the corresponding dimers listed in ii).

The structure of Banf1_Thr2_-DNA was adopted from 2BZF. The structure of Banf1_Met1_-DNA was docked, where the structure of Banf1 monomer was adopted as Chain A in 6UNT and the DNA as Chain B and Chain C in 2BZF. The structures of pSer4 Banf1_Met1_-DNA, pBanf1_Met1_-DNA, pSer4 Banf1_Thr2_-DNA and pBanf1_Thr2_-DNA were generated by docking the representative monomer structure(s) of the corresponding phosphorylated Banf1 to the DNA from 2BZF. HADDOCK 2.4 web server with HADDOCK default settings (https://wenmr.science.uu.nl/haddock2.4/) [40] was used to dock DNA to Banf1, where Banf1 residues within 10 Å from DNA in 2BZF were defined as DNA binding sites.

The structure of WT Banf1_Met1_ dimer was adopted from the Protein Data Bank (PDB ID: 1QCK), whose both monomers contain residue Met1 and their N-terminal conformation is distinct from that of the Banf1 monomer with Met1in the high-resolution crystalised structures (e.g., PDB ID: 1CI4 and PDB ID: 6UNT). We chose this structure as the starting structure of Banf1_Met1_ to validate the reliability of the force fields, as we believe that if the force field is reliable for this project, the representative N-terminal conformations of WT Banf1_Met1_ and pBanf1 from modelling should be consistent with those in the crystalised structures of Banf1_Met1_ (PDBID:6UNT for WT Banf1 and PDBID: 7NDY for pBanf1). The structure of WT Banf1_Thr2_ dimer was obtained by deleting DNA and water in the co-crystalised Banf1-DNA complex (PDB ID: 2BZF). Discovery Studio 2019 [41] was used to build the structures of pSer4 Banf1_Met1_ and pSer4 Banf1_Thr2_ by mutating Ser4 of WT Banf1_Met1_ and WT Banf1_Thr2_ into pSer4, respectively. Similarly, structures of pBanf1_Met1_ and pBanf1_Thr2_ were generated via mutating Thr3 of pSer4 Banf1_Met1_ and pSer4 Banf1_Thr2_ into pThr3 in Discovery Studio 2019 [41]. Structures of unphosphorylated, mono-phosphorylated and di-phosphorylated Banf1_Met1_ and Banf1_Thr2_ monomers were obtained by deleting one monomer of the corresponding Banf1 dimers.

#### Molecular dynamics simulation protocol

Each structure was fully solvated in an octahedral box of TIP3P water [42] with the solute being at least 12 Å away from the edges of the water box. This results in 22 systems of Banf1 protein structures. Periodic boundary conditions were employed. Na+ counter ions were added with tleap in AmberTools 16 [43] to neutralise the net charge of the systems. Sodium chloride (Na+ and Cl-ions) with a concentration of 150 mM were added in all systems to mimic the salt concentration in cells using tleap [43]. In this work, all MD simulations were performed with the PMEMD module (CUDA version) of the AMBER 16 software suite [44, 45] and ff99SB force field for proteins and DNA [46]. Amber S2P and T2P parameters for the phosphoserine and phosphothreonine documented in frcmod.phosaa10 were used for the pSer4 and pThr3 in Banf1, respectively [47, 48], which have been extensively used in previous studies [49–51]. The SHAKE algorithm [52] was applied to restrain the lengths of all covalent bond containing hydrogen atoms, which permits a computational time step of 2 fs. The particle mesh Ewald (PME) summation method [53, 54] was utilised to compute the long-range electrostatic energy, and a cutoff distance of 12 Å was used for the calculation of non-bonded interactions.

In order to obtain a well-equilibrated system for subsequent MD production runs, all systems went through two rounds of geometry optimizations. Firstly, the solute was fixed by a positional restraint of 50 kcal/mol·Å, whereas water molecules were left free for 2000 minimization steps, consisting of 1500 steepest descent steps followed by 500 conjugate gradient steps. Secondly, the geometry of the protein was further optimised with the same number of steepest descent and conjugate gradient steps with both the solute and solvent left free to relax. Then, the optimised system was heated for 500 ps in an NVT ensemble from 100 K to 310 K, where the solute was held fixed with a positional restraint of 50 kcal/mol·Å. Following the NVT annealing, the system was equilibrated by a 500 ps NPT ensemble at a constant pressure of one atmosphere with the Berendsen barostat [55] without applying restraints on either the solute or solvent. Finally, MD production runs were conducted on each equilibrated system without any restraints, where the temperature was held at 310 K by the Langevin thermostat with gamma_ln set to 2 [56], and the pressure was controlled at one atmosphere by the Berendsen barostat [55] and a pressure coupling constant of 1 ps.

Production runs of unphosphorylated Banf1-DNA complexes were carried out for 0.5 μs. Production runs of phosphorylated Banf1-DNA complexes were performed 50 ns. Production runs of WT and mono-phosphorylated Banf1 were performed for 1µs. Simulations of di-phosphorylated Banf1 were performed for 1.5 µs to obtain additional conformational sampling, as the secondary structures of their N-terminal residues were observed to be more dynamic. Systems of Banf1-DNA complexes and Banf1 monomers were all simulated with 3 replicas starting from different atomic velocities. Results for Banf1-DNA complexes were averaged from trajectories of the final 300 ns simulations in the 3 MD simulation replicas. Simulations of Banf1 dimers were simulated with one replica. Results for Banf1 were averaged from the two promoters of the dimer, which is similar to that the systems were simulated with 2 replicas starting from different atomic velocities.

#### Data analysis

Post data processing and analysis were performed with cpptraj in AmberTools 16 [43], VMD 1.9.3 [57], UCSF Chimera 1.15 (https://www.cgl.ucsf.edu/chimera/olddownload.html) [58], GNU bash, and Python 3.8. UCSF Chimera 1.15 [58] was used for visualization and images. For the sake of clarity, one-letter residue code is used to label residues, and some sidechain hydrogens not involved in interactions are hidden in all images of Banf1 conformations and their surface representations. GNUPLOT 5.2 (http://www.gnuplot.info) was used for plotting.

To obtain the representative structures of Banf1 at different phosphorylation status and their occupancies, structural clustering analysis was performed on the MD simulation trajectories of Banf1 structures, with the initial 200 ns trajectories not taken into account. There are different algorithms for clustering conformations of biomolecules. For example, Markovian method can be used to better portray the state change process of the systems of biomolecules, such as our Banf1-DNA complexes [59, 60]. Given that this work focuses on the N-terminal conformational difference of Banf1 induced by its mono- and di-phosphorylation, we chose the hieragglo algorithm in cpptraj in AmberTools 20 [46], which has been popularly used by researchers to cluster the conformation of proteins for simplicity [61, 62]. This clustering method clusters the protein conformations in given frames of corresponding MD trajectories into different clusters based their root-mean-square deviation (RMSD). In this study, the conformation of residue Met1/Thr2 to His7 were used for clustering, with a cutoff value of RMSD set to 3 Å.

The Molecular Mechanics/Generalized Born Surface Area (MM/GBSA) binding free energies between Banf1 and DNA were calculated by the MMPBSA.py script [63] in AmberTools 20 [46]. The MM/GBSA binding free energy (ΔG_bind_) between Banf1 and DNA can be defined as:

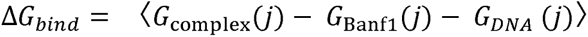

where 〈⋯〉 means averaging over frames from MD trajectory and U denotes Uth snapshot of othe Banf1-DNA complex [64]. Energy of the complex and its individual components can be calculated using the below equations [64]:

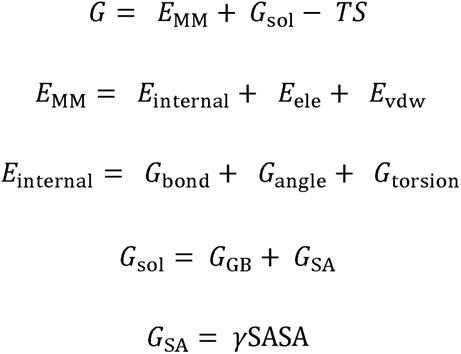

Where E_MM_, G_sol_ and - TΔS represent the gas-phase MM energy, the solvation free energy and the conformational entropy induced by binding, respectively. E_MM_ is composed of the intramolecular bonded energy E_internal_ and non-bonded energy including electrostatic potential E_ele_ and van der Walls energy E_vdw_. G_sol_ is contributed by polar electrostatic solvation energy G_GB_ and the nonpolar nonelectrostatic solvation energy, which can be calculated by GB model and solvent accessible surface area (SASA), respectively.

The occupancies of hydrogen bonds were calculated based on the geometrical criteria where the distance between the acceptor and the donor heavy atoms is less than 4 Å and the donor heavy-atom, donor hydrogen and acceptor bond angle is greater than 120°.

## Results

### Interaction of unphosphorylated Banf1 with DNA

To explore detailed atomic-level interactions between Banf1 and DNA, we performed comprehensive MD simulations on the Banf1_Thr2_-DNA and Banf1_Met1_-DNA complexes. Given that the complex structure of Banf1_Met1_-DNA has not been experimentally characterized, we first validated the reliability of HADDOCK 2.4 [40] in docking DNA to Banf1, and then used it to dock DNA to Banf1_Met1_ for the starting structure of Banf1_Met1_-DNA in MD simulations.

#### HADDOCK2.4 reproduced the binding mode of DNA to Banf1_Thr2_ co-crystalised in 2BZF

To validate the performance of HADDOCK 2.4 [40] in docking DNA to Banf1 protein, DNA from 2BZF was docked to Banf1_Thr2_ extracted from the co-crystalized complex (PDB ID: 2BZF) in the HADDOCK 2.4 server (https://wenmr.science.uu.nl/haddock2.4/). The docked complex adopts the same binding pose as that co-crystalized in 2BZF (Figure 2A), which is reflected by: i) the docked Banf1_Thr2_-DNA overlay with the co-crystalised Banf1_Thr2_-DNA in 2BZF, with a backbone RMSD value of 0.4 Å after fitting the two structures; ii) the docked complex retained all the hydrogen bonds between DNA and Banf1 presented in the co-crystalised 2BZF; and iii) the sidechain directions of Banf1 residues interacting with DNA in the co-crystalised 2BZF are mostly retained in the docked structure with only slight deviation in the direction of sidechains of Lys6, Lys72 and Arg75 as they are long and flexible, which is also reflected in the results from following MD simulation. This demonstrates that HADDOCK 2.4 is reliable for docking DNA to Banf1 protein.

**Figure 2.**
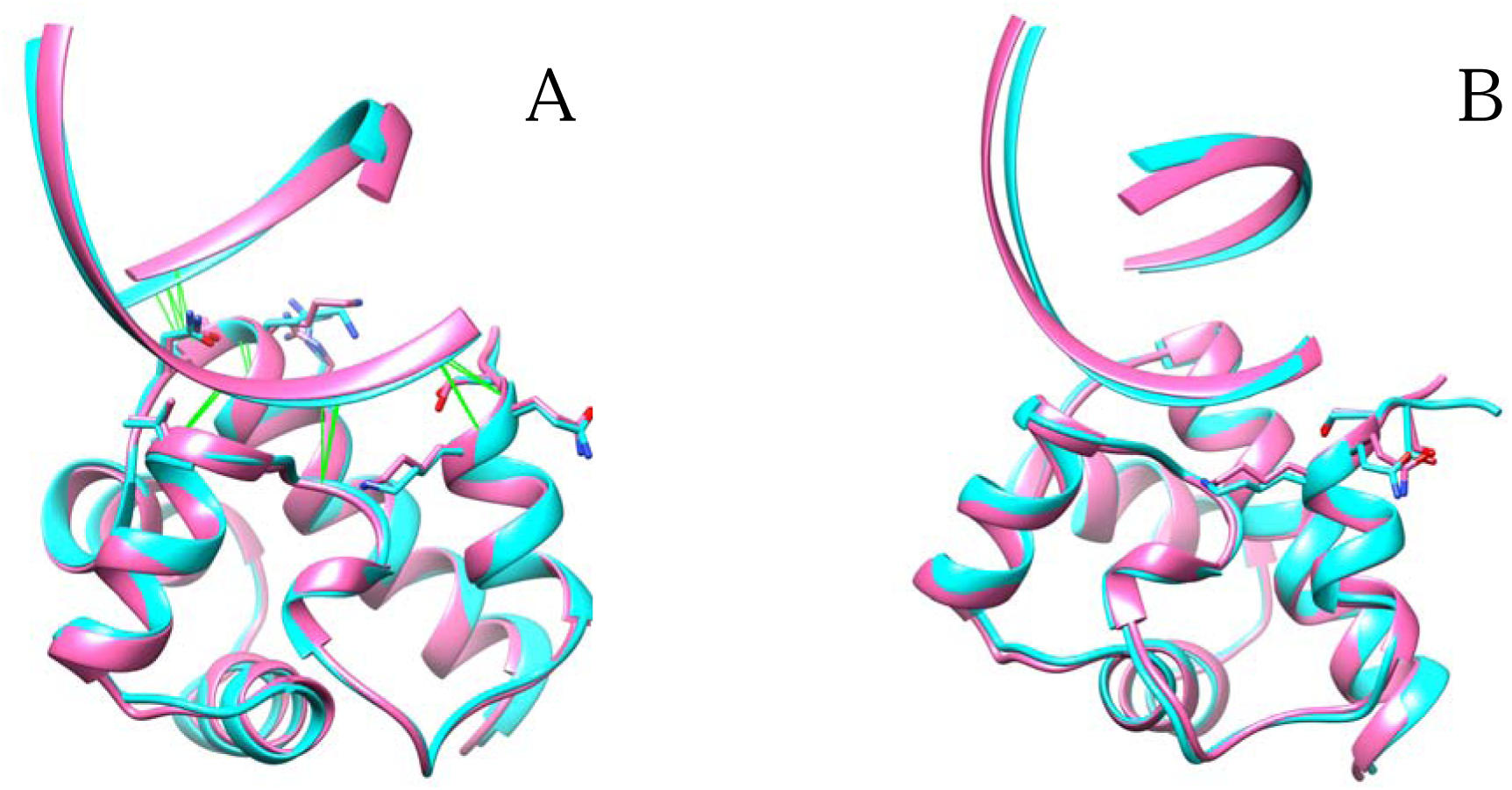
A Overlay of the docked Banf1_Thr2_-DNA complex from HADDOCK 2.4 (cyan) to the co-crystalised complex (PDB ID: 2BZF, hot pink). B Superimposed complex structure of the Banf1_Thr2_-DNA (hot pink) and docked Banf1_Met1_-DNA (cyan). Sidechain of residues 3-6 in Banf1 of both complexes are displayed without hydrogen atoms for the sake of clarity.

#### Binding mode of DNA to Banf1_Met1_ predicted by HADDOCK 2.4

Figure 2B shows the docked structure of the Banf1_Met1_-DNA complex, superimposed with the co-crystalised Banf1_Thr2_-DNA complex. As indicated in Figure 2B, the binding mode of DNA to Banf1_Met1_ in the docked complex is quite close to the mode co-crystalised in 2BZF; the Banf1_Met1_-DNA approximately overlaps with the co-crystalised Banf1_Thr2_-DNA complex, with an RMSD value of 0.1 Å. Similarly, the docked Banf1_Met1_-DNA retains all Banf1-DNA inter-molecular hydrogen bonds presented in the co-crystalised complex. Notably, Met1 is not involved in interacting with DNA in the Banf1_Met1_-DNA complex, which is consistent with previous docking results from ZDOCK [65].

#### Stability and convergence of MD simulations

The binding poses of Banf1_Thr2_-DNA and Banf1_Met1_-DNA remained stable in the three parallel replicas of MD simulations starting from different atomic velocities. This is supported by the low and stable RMSD time evolution of Banf1 in Banf1_Thr2_-DNA (green) and Banf1_Met1_-DNA (red) during the 500-ns-long MD simulations (Figure 3A). The stability of the Banf1-DNA complexes in MD simulations is also reflected in the low RMSF values of Banf1 residues shown in Figure 3B. Specifically, apart from the N and C terminal, Banf1 protein is stable when bound to DNA, with most residue has an RMSF below 1 Å. This aligns with previous X-ray crystallographic experimental results that binding to DNA does not change Banf1’s structural conformation [11]. The N-terminal region containing Met1, Thr2, Thr3 and Ser4, are very flexible, as reflected by their much higher RMSF (Figure 3B). This is in consistent with the recent and earlier NMR results which unravelled that the N-terminal region of Banf1 is very flexible in solution [17, 66, 67].

**Figure 3.**
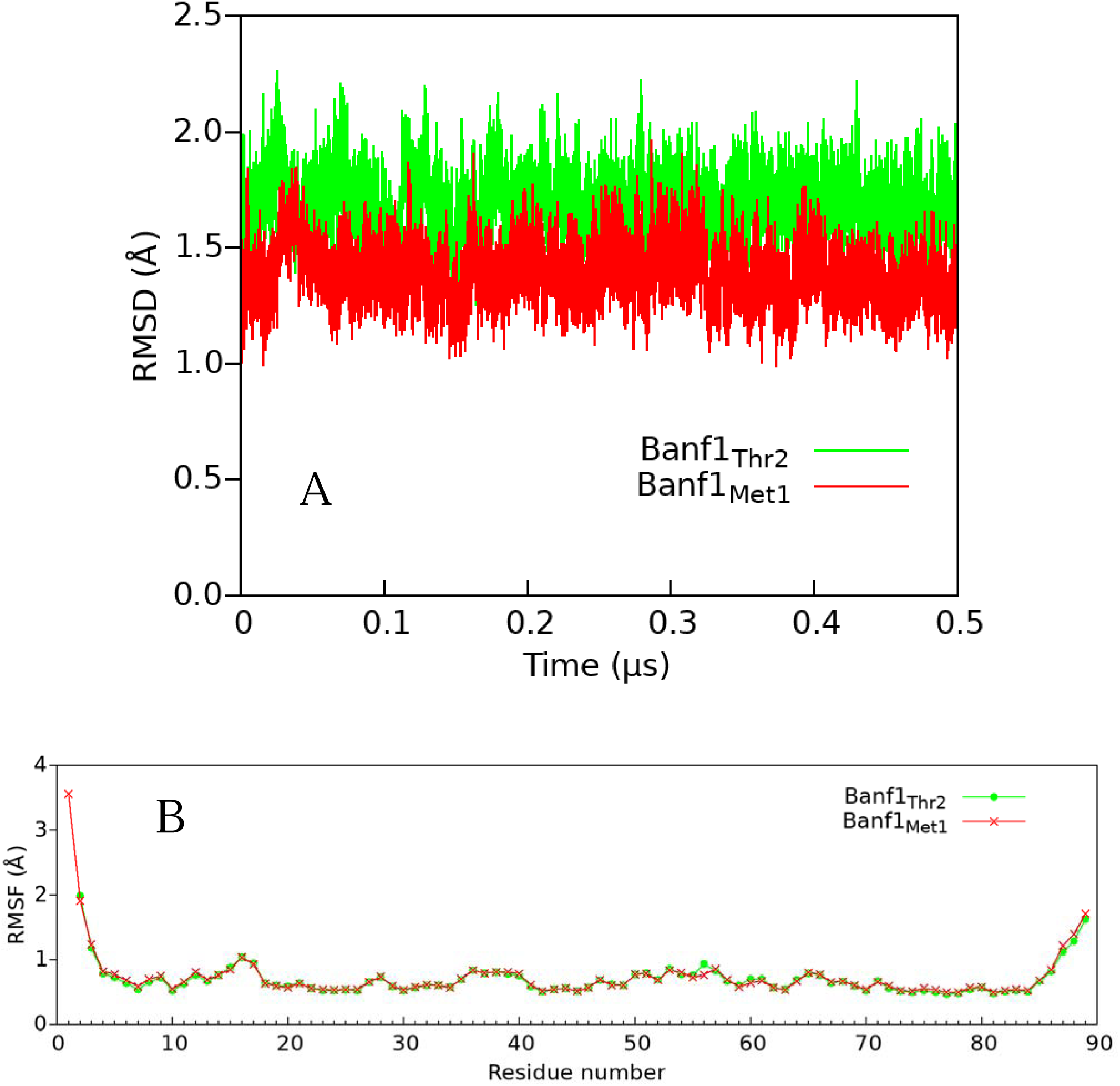
Stability and convergence of MD simulations of Banf1-DNA complexes. A RMSD variations with respect to time of Banf1_Thr2_-DNA (green) and Banf1_Met1_-DNA (red). B RMSF variations with respect to residues of Banf1_Thr2_ (green) in the Banf1_Thr2_-DNA complex and Banf1_Met1_ (red) in the Banf1_Met1_-DNA complex.

#### Binding energy between Banf1 and DNA

To further explore if containing Met1 in Banf1 makes any difference in the Banf1-DNA binding energy, we calculated the total binding energies and their components for the Banf1_Thr2_-DNA and Banf1_Met1_-DNA complexes (Table 1). As shown in Table 1, Banf1_Thr2_ and Banf1_Met1_ have a similar DNA-binding free energy (ΔG_bind_), where Banf1_Thr2_ binds to DNA with an approximately 3% higher affinity than Banf1_Met1_. This agrees with the our and published docking results [65] that containing Met1 at the N-terminus of Banf1 does not affect its DNA binding, and is also in line with the fact that both Banf1 with and without Met1 function well in cells.

**Table 1.** MMGBSA binding energies of Banf1 and DNA averaged from 3 independent MD simulation replicas starting from different atomic velocities.

| Energy | $\Delta E_{\text{vdw}}$ | $\Delta E_{\text{ele}}$ | $\Delta G_{\text{GB}}$ | $\Delta G_{\text{SA}}$ | $\Delta G_{\text{bind}}$ |
| --- | --- | --- | --- | --- | --- |
| Banf1<sub>Met1</sub>-DNA | -59.6±4.6 | -742±93.6 | -727.9±88.6 | -7.1±0.5 | -80.7±10.9 |
| Banf1<sub>Thr2</sub>-DNA | -59.1±4.4 | 706.5±84.5 | 695.3±80 | -6.8±0.5 | -77.1±9.7 |

To identify which and to what extent residues in Banf1-DAN complexes contribute to the binding, we calculated the backbone and sidechain energy contribution of all residues in the Banf1_Thr2_-DNA and Banf1_Met1_-DNA complexes (Figure 4). As illustrated in Figure 4, residues involved in binding are consistent and their energy contribution, including both the backbone and the sidechain, are comparable in the two Banf1-DNA complexes. Notably, energy contribution of the N terminal residues 1-3 in Banf1 are almost negligible for its DNA binding. In consistent with the X-ray crystallography characterisation [11], the backbone of Gln5, Lys6, Gly25, Gly27, Val29, Leu30, and Ala71, and the sidechain of Lys6, Val29 and Arg85 in Banf1 contribute significantly to its DNA binding, regardless of whether or not the protein has Met1 (Figure 4). Similarly, the sidechain of Ser4 in Banf1 contributes to its DNA binding with a consistent amount of ∼-0.85 kcal/mol, consistent with the fact that it forms a water bridge with the phosphate of DNA nucleotide DA6 [11]. Notably, the sidechain of Lys6 in Banf1 contributes significantly to its DNA binding (Figure 4), in line with previous experimental studies that K6A Banf1 mutant showed indetectable DNA binding property [68]. These results further demonstrate that our modelling is capable of characterising Banf1-DNA interactions.

**Figure 4.**
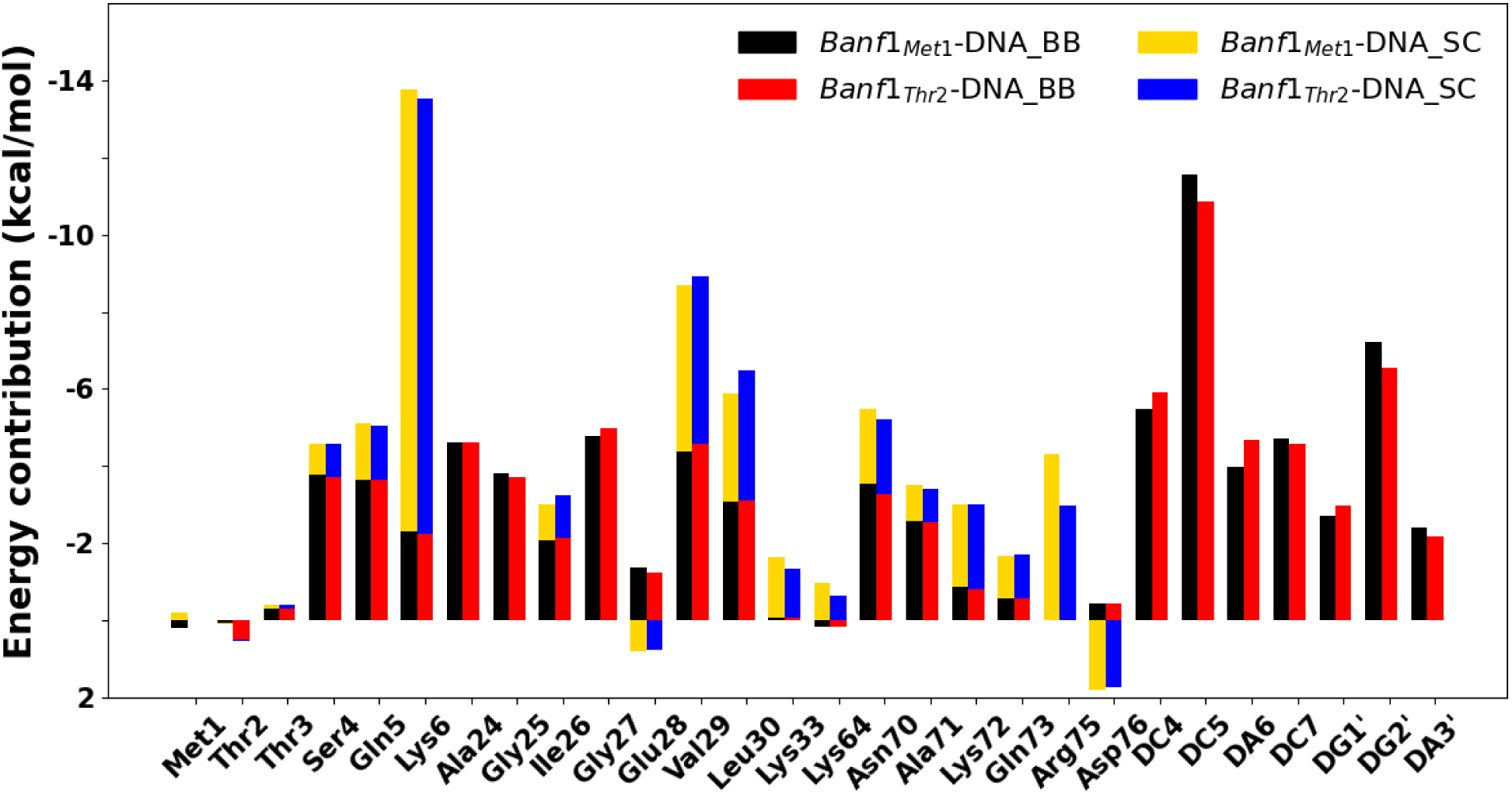
Interaction energies between DNA and the backbone and sidechain of some important Banf1 residues in Banf1_Thr2_-DNA and Banf1_Met1_-DNA. In the legends, BB is short for backbone and SC for sidechain. For the sake of clarity, only N-terminal residues and residues that contribute more than −1 kcal/mol (total energy) are displayed. Negative energy contribution suggests the sidechain/backbone contributes to the binding, whereas positive energy contribution indicates the sidechain/backbone reduces to the binding. For the sake of clarity, the energy contribution of the sidechain of DNA nucleotides are not shown, given that Banf1 residues mostly interact with DNA backbone.

Apart from the abovementioned 9 residues characterised by X-ray crystallography that stabilise the Banf1-DNA complex, there are some other intermolecular interactions that contribute to the Banf1-DNA binding. For example, the amide nitrogen of Ser4, Ala24, Ile26, Glu28, Asn70 and Lys72 in Banf1 can form electrostatic interactions or transient hydrogen bonds (Glu28) with DNA phosphates and contribute remarkably to the binding energy, due to their close distance (within 5 Å). Moreover, the sidechain of polar residues such as Gln5, Lys33, Lys64, Lys72 and Gln73 in Banf1 also contribute to its DNA-binding, because they can form transient direct hydrogen bonds and/or water-mediated hydrogen bonds with DNA nucleotides as discussed in the following section. In contrast, the sidechains of non-polar residues such as Ile26, Leu30 and Ala71 in Banf1 form Van der Waals interactions with the ribose of DNA nucleotides. Surprisingly, the sidechain of Glu28 and Asp76 in Banf1 impair its DNA binding capability, which has not been reported in the literature.

#### Banf1-DNA intermolecular Hydrogen bonding interactions

Inter-molecular hydrogen bonding interaction is the main contributor of Banf1-DNA binding energy [11]. To understand the atomic basis of the above-presented residue-level energy contribution to the Banf1-DNA binding, we investigated the direct (Table 2) and water-mediated indirect (Table 3) hydrogen bonds between Banf1 and DNA. As illustrated in Table 2, all of the 9 hydrogen bonds characterised by X-ray crystallography [11] were highly frequently kept in both Banf1-DNA complexes. This demonstrates the reliability of the molecular modelling protocol, i.e., docking in HADDOCK 2.4 followed by MD simulations, in investigating the interaction of Banf1 with DNA. Interestingly, in the two complexes there are some less frequent, but highly consistent hydrogen bonds between DNA backbone and Banf1 polar residues. Specifically, the amide nitrogen of Gln28 forms a hydrogen bond with the phosphate of nucleotide DC5 with an occupancy of 20-31%. With a similar frequency, the ε-amino group of Lys33 and Lys64 forms hydrogen bonding interaction with the phosphate of nucleotide DC4 and DG1’, respectively. In addition, there is a hydrogen bond with an occupancy of 24-26% between the amide group of Gln73 and the oxygen atom in the ribose (O4’) of nucleotide DG4’. The newly found hydrogen bonds and their occupancies are comparable in the two complexes and are consistent in all the three parallel simulations carried out on each of both complexes.

**Table 2.** Intermolecular hydrogen bonds between Banf1 and DNA with an occupancy higher than 20%. Data are averaged from 3 independent MD simulation replicas starting from different atomic velocities.

| Donor | DonorH | Acceptor | Occupancy (%) |  |
| --- | --- | --- | --- | --- |
|  |  |  | Banf1<sub>Met1</sub>-DNA | Banf1<sub>Thr2</sub>-DNA |
| Leu30@N | Leu30@H | DC5@O1P | 100 | 100 |
| Gly25@N | Gly25@H | DA6@O1P | 100 | 99 |
| Gln5@N | Gln5@H | DC7@O1P | 96 | 95 |
| Val29@N | Val29@H | DC5@O2P | 95 | 89 |
| Lys6@NZ | Lys6@HZ1/2/3 | DA6@O2P | 94 | 93 |
| Ala71@N | Ala71@H | DG1'@O1P | 93 | 93 |
| Lys6@N | Lys6@H | DC7@O2P | 91 | 87 |
| Gly27@N | Gly27@H | DC5@O1P | 79 | 93 |
| Val29@N | Val29@H | DC5@O1P | 71 | 86 |
| Arg75@NH1/2 | Arg75@HH12/22 | DG2'@O1P | 100 | 80 |
| Glu28@N | Glu28@H | DC5@O2P | 31 | 20 |
| Lys33@NZ | Lys33@HZ1/2/3 | DC4@O1P | 30 | 21 |
| Lys64@NZ | Lys64@HZ1/2/3 | DG1'@O1P | 21 | 15 |
| Gln73@NE2 | Gln73@HE21 | DC4@O4' | 24 | 26 |

**Table 3.**
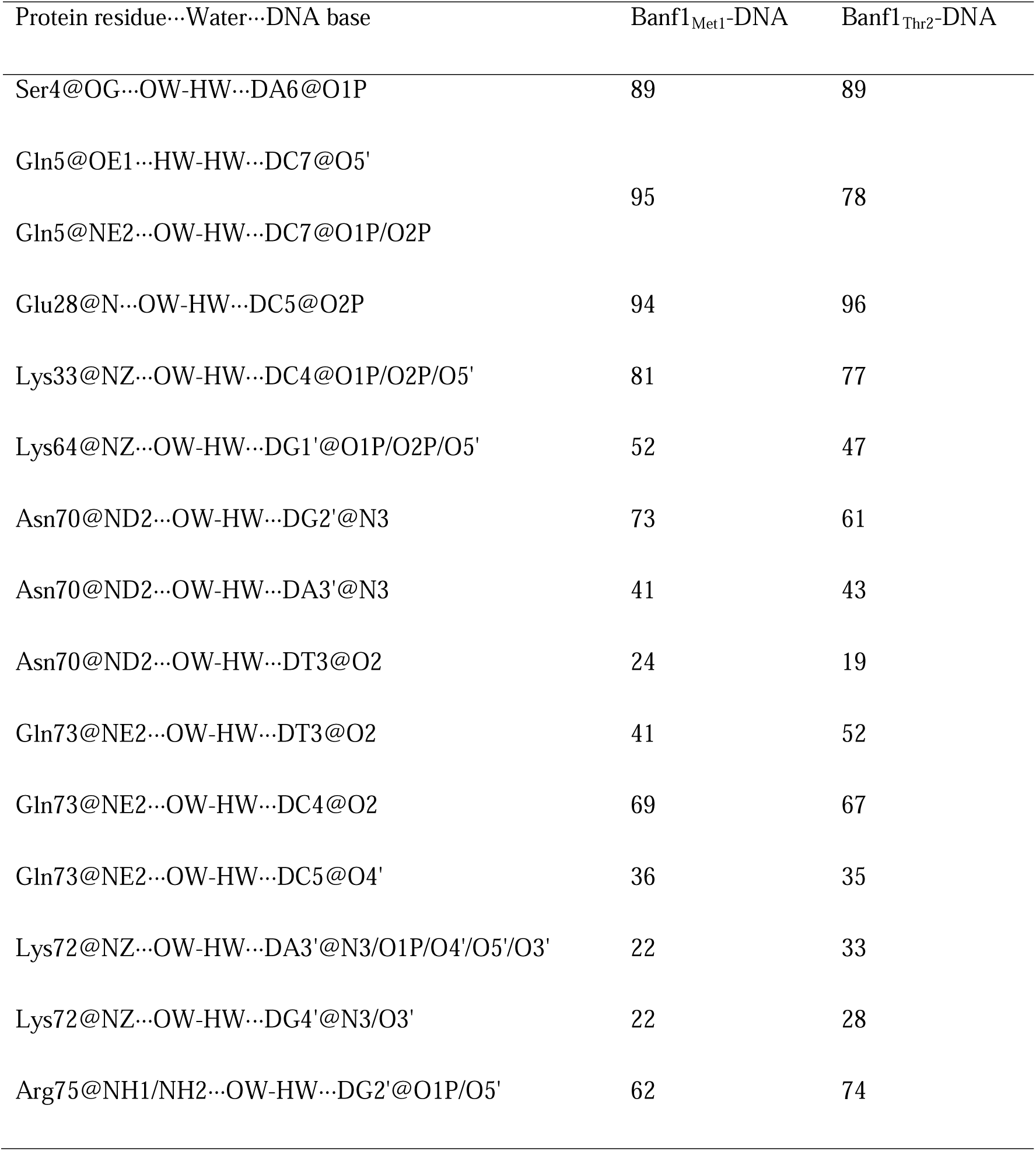
Water bridges between Banf1 and DNA with an occupancy higher than 20%. Data are averaged from 3 independent MD simulation replicas starting from different atomic velocities.

| Protein residue...Water...DNA base | Banf1<sub>Met1</sub>-DNA | Banf1<sub>Thr2</sub>-DNA |
| --- | --- | --- |
| Ser4@OG...OW-HW...DA6@O1P | 89 | 89 |
| Gln5@OE1...HW-HW...DC7@O5' | 95 | 78 |
| Gln5@NE2...OW-HW...DC7@O1P/O2P |  |  |
| Glu28@N...OW-HW...DC5@O2P | 94 | 96 |
| Lys33@NZ...OW-HW...DC4@O1P/O2P/O5' | 81 | 77 |
| Lys64@NZ...OW-HW...DG1'@O1P/O2P/O5' | 52 | 47 |
| Asn70@ND2...OW-HW...DG2'@N3 | 73 | 61 |
| Asn70@ND2...OW-HW...DA3'@N3 | 41 | 43 |
| Asn70@ND2...OW-HW...DT3@O2 | 24 | 19 |
| Gln73@NE2...OW-HW...DT3@O2 | 41 | 52 |
| Gln73@NE2...OW-HW...DC4@O2 | 69 | 67 |
| Gln73@NE2...OW-HW...DC5@O4' | 36 | 35 |
| Lys72@NZ...OW-HW...DA3'@N3/O1P/O4'/O5'/O3' | 22 | 33 |
| Lys72@NZ...OW-HW...DG4'@N3/O3' | 22 | 28 |
| Arg75@NH1/NH2...OW-HW...DG2'@O1P/O5' | 62 | 74 |

The occupancies of water bridges between Banf1 and DNA are comparable in the two complexes (Table 3). In consistent with the characterisation of X-ray crystallography, there is a water bridge with an occupancy of around 90% between the sidechain of Ser4 and the phosphate of nucleotide DA6. Here, we also found some new frequent water-mediated hydrogen bonds between Banf1 polar residues and the nitrogen and oxygen atoms in DNA nucleotides (Table 3). Specifically, the sidechain of Gln5 forms a water bridge with the backbone of nucleotide DC7 with a high frequency of 78-95%. Interestingly, apart from forming a direct hydrogen bond with nucleotide DC5 via the amide nitrogen, Glu28 frequently forms a water bridge with nucleotide DC5 via its amide nitrogen and γ carboxyl group. Furthermore, Banf1 positively charged residues with long a sidechain located close to the DNA binding surface form frequent water bridges with DNA bases. For example, the ε-amino group of Lys33, Lys64 and the guanidine group of Arg75 form a water bridge with the backbone of nucleotide DC4, DG1’ and DG2’ with an occupancy of ∼80%, ∼50% and ∼70%, respectively. Interestingly, sidechain of Lys72 under DNA strand 2 (DG1’-DG7’) that interact with Banf1’s pseudo HhH motif, form water bridges with both the sidechain and backbone of nucleotide DA3’ and DG4’. In addition, the amide group of Asn70 and Gln73 also interact with multiple DNA nucleotides. Particularly, the sidechain of Asn70, located under the sidechains of both DNA strands, form water-mediated hydrogen bonds with the backbone of two adjacent nucleotides (DG2’ and DA3’) in DNA strand 2 and one nucleotide (DT3) in DNA strand 1 (DC1-Dc7). Similarly, the amide group of Gln73 interacts with three adjacent nucleotides (DT3, DC4 and DC5) in DNA strand 1 via water-mediated hydrogen bonds. We note that the water bridges between Banf1 and the sidechains of DNA nucleotides may vary with the nucleotide sequence of the DNA. These results demonstrate that various Banf1 polar residues form frequent water-mediated hydrogen bonding interactions with DNA nucleotides, which contribute to the stabilisation of the Banf1-DNA complex.

### Mechanisms of N-terminal phosphorylation in Banf1 reducing its DNA binding capability

To investigate the dynamics of the N-terminal secondary structure of di-phosphorylated Banf1 (hereafter denoted as pBanf1) and how di and mono-phosphorylation of Banf1 (hereafter denoted as pSer4 Banf1) impair the above presented atomic bases of Banf1-DNA binding and consequently eliminate its DNA binding capability, we performed systematic and comprehensive MD simulations to explore if and how mono- and di-phosphorylation impact the N-terminal secondary structure of Banf1_Met1_ and Banf1_Thr2_ as well as their DNA binding surfaces.

#### Phosphorylation does not induce major conformational changes in residues 7-89 of Banf1

In line with the X-ray diffraction experiments [17], our MD simulations confirmed that phosphorylation does not induce any distinct conformational changes in residues 7-89 of Banf1. This is supported by the low RMSD in the backbone of residues 7-89 in pSer4 Banf1 and pBanf1 displayed in Figure 5 for Banf1_Thr2_ and in Supplementary Figure S1 for Banf1_Met1_. As illustrated in Figure 5 and Figure S1, RMSD of residues 7-89 in WT Banf1, pSer4 Banf1 and pBanf1 are stable at ∼1 Å throughout the simulations, which is independent of whether Banf1 contains Met1 on the N-terminus. The stability of residues 7-89 in Banf1 with different phosphorylation status is also clearly reflected in their low RMSF in Figure 6 and Figure S2, as well as in their representative conformations (Figure 7 and S3) determined from MD simulations to be discussed in the following section.

**Figure 5.**
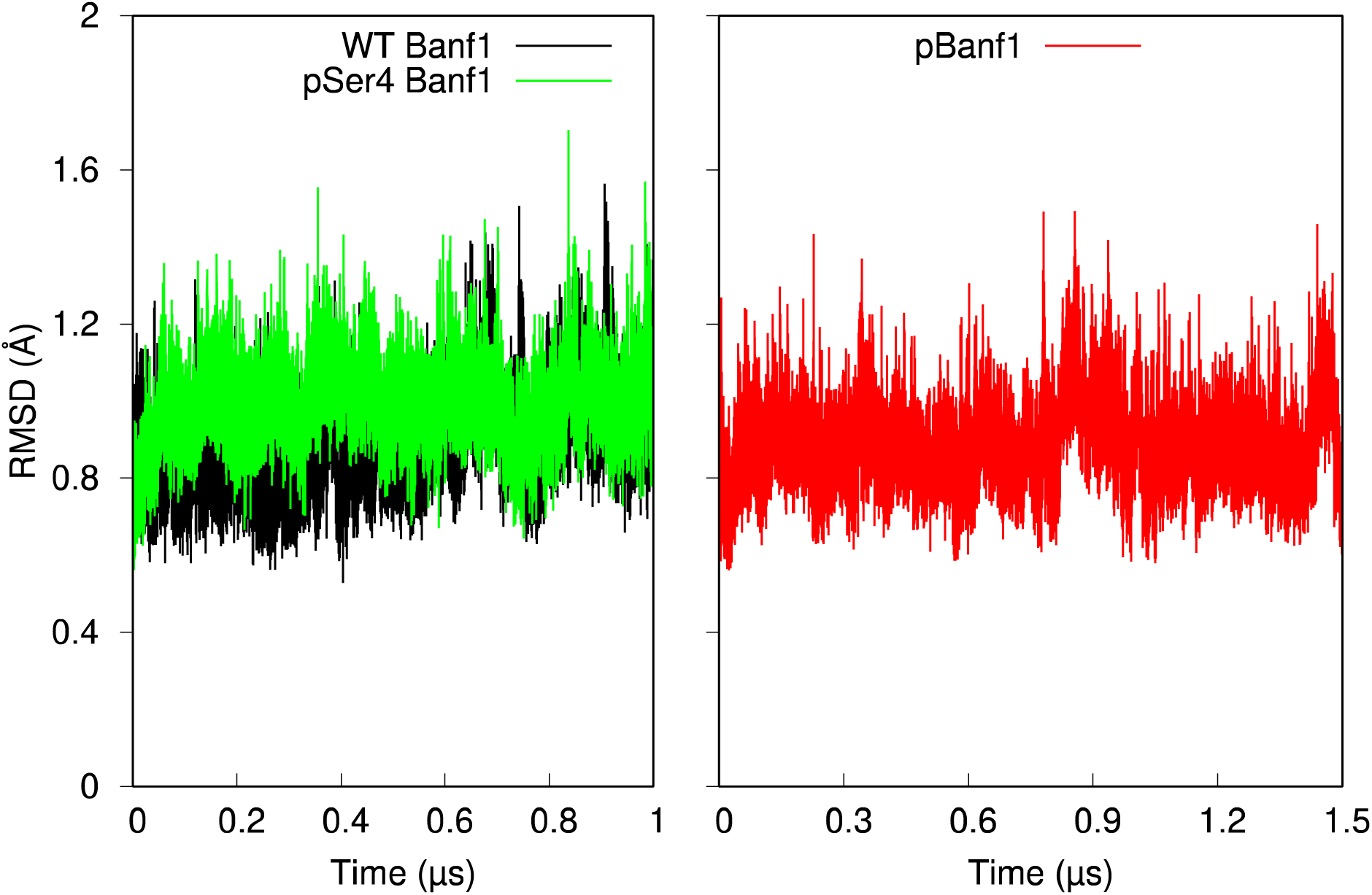
RMSD variations with respect to time of residues 7-89 in Banf1_Thr2_, where WT Banf1 is for unphosphorylated Banf1_Thr2_, pSer4 Banf1 for mono-phosphorylated Banf1_Thr2_, and pBanf1 for di-phosphorylated Banf1_Thr2._

**Figure 6.**
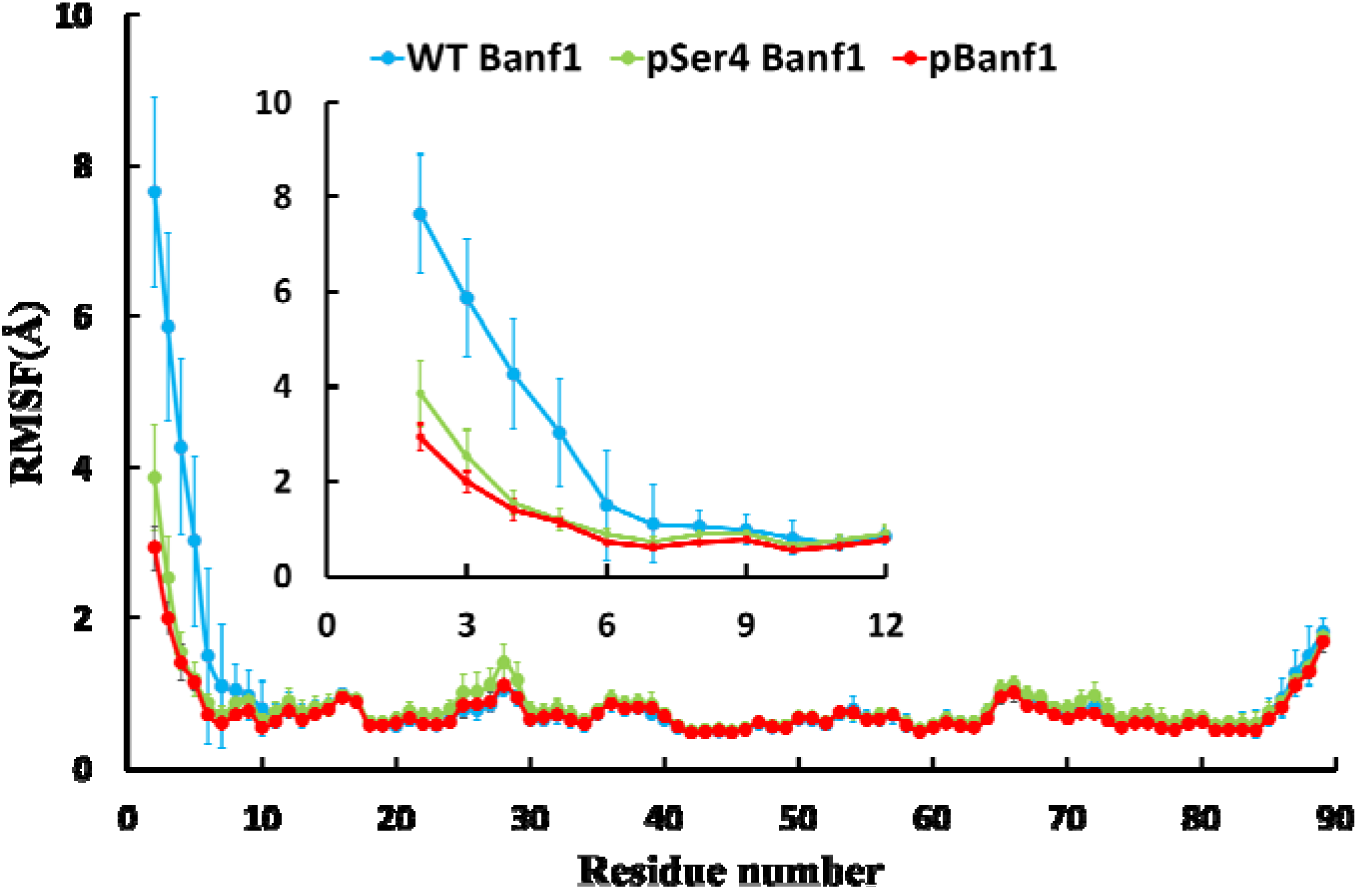
RMSF and its standard deviation of residues 2-89 in Banf1_Thr2_ and its zoom in view incorporating the N-terminal residues 2-12 of Banf1_Thr2_. The RMSF values were averaged from the RMSF of the two Banf1_Thr2_ promoters in the simulation of Banf1_Thr2_ dimer and the RMSF of the Banf1_Thr2_ monomer in the three simulation replicas of Banf1_Thr2_ monomer. Trajectories from the initial 100 ns simulations of unphosphorylated Banf1_Thr2_ was discarded and the trajectories of simulations before the mono- and di-phosphorylated Banf1_Thr2_ form an additional N-terminal helix were ignored.

**Figure 7.**
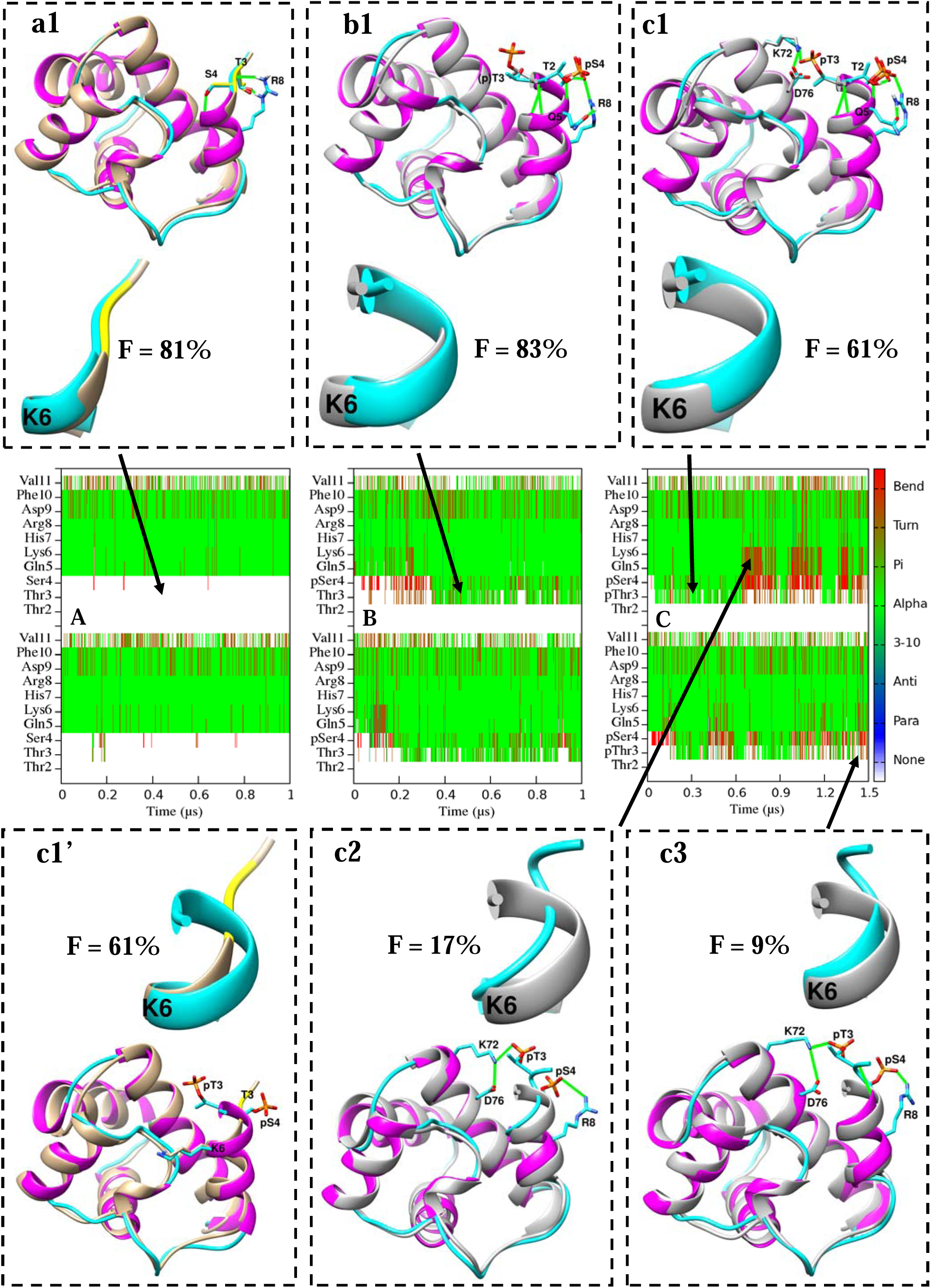
Secondary structure maps of the N-terminal residues 2-11 in two protomers of Banf1_Thr2_ dimer and the corresponding Banf1_Thr2_ representative conformations superimposed with reference structures. A, B, and C are the secondary structure maps of unphosphorylated Banf1_Thr2_, mono-phosphorylated Banf1_Thr2_ and di-phosphorylated Banf1_Thr2_, respectively. a1 and b1 are the representative conformation of unphosphorylated and mono-phosphorylated Banf1_Thr2_, respectively. c1, c2 and c3 are the representative conformations of di-phosphorylated Banf1_Thr2_. c1’ displays the same representative conformation as c1 but with a different reference structure. Zoom-in view of the N-terminal residues 2-7 of each representative structure is displayed in ribbon. Occupancies (F) of each representative conformation of Banf1 are also displayed accordingly. Conformations from MD are coloured by secondary structure, with helical structure in magenta and coil in cyan. Reference structures in a1 and c1’ coloured in tan are adopted from chain A of 2BZF, with the phosphorylation sites of Thr3 and Ser4 highlighted in yellow. Reference structures for other representative conformations coloured in gray are chain A extracted from 7NDY. For clarity, sidechains of pSer4 and pThr3 in the reference structures in c2 and c3, and the phosphate group of pThr3 in the reference structure in c1 are not displayed. Hydrogen bonds and salt bridges are shown as green lines. Sidechains in reference structures are coloured in the same colour as their chain, whereas sidechains in MD representative structures are coloured in cyan. Oxygen, nitrogen, hydrogen and phosphorus atoms are coloured in red, blue, white and gold respectively.

#### Phosphorylation induces distinct local conformational changes in the N-terminal region of Banf1

As illustrated in Figure 1B and 1C, the N-terminal residues Ser4, Gln5 and Lys6 in Banf1 play an essential role in achieving its binding to DNA [11], indicating that conformational changes in the N-terminal region of Banf1 may alter its DNA binding capability. Hence, this section investigates the impacts of mono- and di-phosphorylation of Banf1 on its N-terminal secondary structure.

Figure S2 and Figure 6 display the RMSF and its standard deviation of the backbone of residues 1/2-89 and its zoom in view incorporating the N-terminal residues 1/2-12 of Banf1_Met1_ Banf1_Thr2_, respectively. As illustrated in Figure S2 and Figure 6, phosphorylation of Banf1 reduces the flexibility of its N-terminal residues. Figure S3 and Figure 7 display the secondary structure variations with respect to time of the N-terminal residues from Met1/Thr2 to Val11 in both protomers of unphosphorylated, mono-phosphorylated and di-phosphorylated Banf1_Met1_ and Banf1_Thr2_ dimer, as well as the representative Banf1 conformations superimposed with the reference structures and the corresponding occupancies. As shown in Figure 7 and S3, the representative conformational structure(s) adopted by the N-terminal residues of Banf1’s two protomers during MD simulations are consistent, although the N-terminal conformation of each protomer in the Banf1 dimer at the same time point might be different due to its flexibility [17, 66]. This aligns with the dynamic nature of proteins in solution. In agreement with the NMR results [17, 66], MD simulations indicate that the N-terminal residues 1-4 of unphosphorylated Banf1 (hereafter denoted as WT Banf1) are highly flexible. Notably, MD simulations of WT Banf1 frequently reproduce the conformations of Banf1 that were previously characterised by X-ray crystallography and NMR (Figure 7 and S3). Specifically, the representative conformation (with an occupancy of 81%) of WT Banf1_Thr2_ suggested by MD overlays with the Banf1_Thr2_ structure co-crystalised in 2BZF (Figure 7a1), with an RMSD of 0.8 Å when fitting them. Similarly, the MD representative conformation (with an occupancy of 52%) of Banf1_Met1_ overlays the structure of Banf1_Met1_ characterised by NMR in 6UNT (Figure S3a2), with an RMSD of 0.8 Å when superimposing them. These results demonstrate that the MD force fields applied in this study are capable of capturing the correct secondary structure of Banf1_Met1_ and Banf1_Thr2,_ including their flexible N-terminal region.

As displayed in the secondary structure maps in Figure 7 and S3, both mono- and di-phosphorylation of Banf1 induce distinct secondary structure changes in residues near the phosphorylation site(s); residues 3 and 4 that either did not have a secondary structure in WT Banf1_Thr2_ or sometimes showed as bend structure at Thr3 in WT Banf1_Met1_, adopted a secondary structure in phosphorylated Banf1 during most of the simulation time. Specifically, the secondary structures of residues 3 and 4 in pBanf1 are dynamic and are transiently occupied as helical, turn or bend structure as illustrated in Figure 7C for pBanf1_Thr2_ and Figure S3C for pBanf1_Met1_. Therefore, both pBanf1_Thr2_ and pBanf1_Met1_ have three representative conformations displayed in Figure 7 and S3, respectively. Consistent with the corresponding small and stable RMSD and RMSF of residues 7-89 discussed in the previous section, di-phosphorylation does not induce major structural changes in residues 7-89 of Banf1. Particularly, residues 7-89 in the three representative conformations of pBanf1_Thr2_ displayed in Figure 7c1, 7c2 and 7c3 have an RMSD of 0.9 Å, 0.7 Å 0.8 Å compared to those in Banf1 co-crystalised in 2BZF. This agrees with the findings from X-ray diffraction experiments that Banf1_Thr2_ does not undergo major structural changes after di-phosphorylation [17]. However, there are significant conformational changes in the N-terminal regions in all the three representative conformations of pBanf1_Thr2_ compared to that in the WT Banf1. Particularly, the representative conformation of pBanf1_Thr2_ in Figure 7c1 (with an occupancy of 61%) overlays with the pBanf1_Thr2_ structure co-crystalised with emerin (PDB ID: 7NDY), with an RMSD of 1.0 Å for all backbone atoms. Additionally, the representative conformation in Figure 7c2 (with an occupancy of 17%) is consistent with the pBanf1_Thr2_ structure characterised by X-ray crystallography for the Banf1 dimer whose N-terminal α1 helix is significantly shortened to residues 7-11. These results further demonstrate the reliability of the modelling. The representative conformations of pBanf1_Met1_ are similar as those of pBanf1_Thr2_ but with different occupancies, as illustrated in Figure S3c1, S3c2 and S3c3. Unlike pBanf1 whose N-terminal secondary structure is relatively dynamic, residues 3 and 4 in pSer4 Banf1 mostly adopt helical structure, and their helical structures are slightly more stable when Met1 is not cleaved (Figure 7B, 7b1, S3B and S3b1). This results in a unique representative conformation adopted by pSer4 Banf1 with an occupancy of 83% for pSer4 Banf1_Thr2_ (Figure 7b1) and 94% for pSer4 Banf1_Met1_ (Figure S3b1), which is similar to one of the representative conformations of pBanf1 displayed in Figure 7c1 and S3c1. Strikingly, pSer4 Banf1_Met1_ and pSer4 Banf1_Thr2_ adopt the same N-terminal secondary structure as that of pBanf1 co-characterised in 7NDY, with a backbone RMSD of 0.8 Å. Given that the electrostatic interactions between both phosphorylated residues (pThr3 and pSer4) and the charged residues Lys72 and Arg75 in the nearby α6 helix were speculated by Marcelot et al. to be essential for restricting the position of the N-terminal region of pBanf1, it would be interesting to reveal the atomic-level interactions that stabilise the same N-terminal secondary structure of pSer4 Banf1 and pBanf1 with a different number of phosphorylated residues.

Table 4 presents occupancies of the secondary structures adopted by residues 2-6 for Banf1_Thr2_ and by residues 1-6 for Banf1_Met1_. Clearly, residues 3-4 in WT Banf1_Thr2_ protomers (Thr3 and Ser4) do not have a secondary structure throughout the simulation (Table 4, Figure 7A and 7a1). In contrast, Thr3 is presented as a helical structure and turn structure with a fraction of 61% and 19%, and pSer4 with a fraction of 78% and 18% in pSer4 Banf1_Thr2_ during the simulation, respectively. Further phosphorylation at Thr3 of pSer4 Banf1_Thr2_ reduces the helical occupancies of residue 3 and 4 to 38% and 49% in the simulation of pBanf1_Thr2_, respectively. Similarly, Thr2, which did not have any secondary structure in WT Banf1_Met1_, presented as a helical structure for a propensity of 8% in pSer4 Banf1_Met1_ and as a helical structure and turn structure with a fraction of 10% and 19% respectively in pBanf1_Met1_. Additionally, Thr3 was shown as bend structure and no structure in WT Banf1_Met1_ for an occupancy of 42% and 57% respectively, but was mostly occupied as helical structure (86%) and turn structure (12%) in pSer4 Banf1_Met1_. Further phosphorylation at Thr3 of pSer4 Banf1_Met1_ significantly reduces the helical occupancy of residue 3 to 46% and increases its turn structure and bend structure ratio to 35% and 18% respectively. In addition, residue 4, which did not have any secondary structure in WT Banf1_Met1_, displayed as a helical structure and turn structure with a propensity of 88% and 11% in pSer4 Banf1_Met1_. Further phosphorylation at Thr3 of pSer4 Banf1_Met1_ results in a lower propensity of helical structure (52%) and a higher occupancy of turn structure (34%) and bend structure (10%) for pSer4. Notably, residues Gln5 and Lys6 are mostly helical structure in WT and pSer4 Banf1, regardless of whether the protein has Met1 or not. However, di-phosphorylation of Banf1 significantly reduces the helical structure occupancies of Gln5 and Lys6 to around 77% from ∼98% in pSer4 Banf1, demonstrating that the N-terminal α1 helix is frequently shortened as compared to WT Banf1. The corresponding representative conformations with shortened N-terminal α1 helix are displayed in Figure 7c2 for Banf1_Thr2_ and Figure S3c2 for Banf1_Met1_. This is consistent with the recent observation by X-ray crystallography that Banf1_Thr2_ N-terminal α1 helix is significantly shortened to comprise residues 7-11 after di-phosphorylation.

**Table 4.** Occupancies of secondary structures for the N-terminal residues in unphosphorylated, mono-phosphorylated and di-phosphorylated Banf1_Thr2_ and Banf1_Met1_. Occupancies of Pi helical structure, anti-parallel beta structure and parallel beta structure are zero and not shown here. Helical structure is mostly alpha helical structure with a very minor ratio of 3-10 helix for several residues. Apart from the total occupancy of helical, turn and bend structures, the remaining occupancy for a residue corresponds to no secondary structure. For example, Thr3 does not have secondary structure in WT Banf1_Met1_ for (100-42-1)% of the simulation time. The occupancies were averaged from two protomers of the dimer. Res3 is Thr3 in WT and pSer4 Banf1, and is pThr3 in pBanf1. Res4 is Ser4 in WT Banf1, and is pSer4 in mono- and di-phosphorylated Banf1. For the sake of clarity, WT represents WT Banf1 and pSer4 is pSer4 Banf1.

| Banf1_Residue | Helical (%) |  |  | Turn (%) |  |  | Bend (%) |  |  |
| --- | --- | --- | --- | --- | --- | --- | --- | --- | --- |
|  | WT | pSer4 | pBanf1 | WT | pSer4 | pBanf1 | WT | pSer4 | pBanf1 |
| Banf1<sub>Thr2</sub>_Thr2 | 0 | 0 | 0 | 0 | 0 | 0 | 0 | 0 | 0 |
| Banf1<sub>Thr2</sub>_Res3 | 0 | 61 | 38 | 0 | 19 | 21 | 0 | 0 | 0 |
| Banf1<sub>Thr2</sub>_Res4 | 1 | 78 | 49 | 1 | 18 | 26 | 2 | 3 | 14 |
| Banf1<sub>Thr2</sub>_Gln5 | 95 | 98 | 76 | 4 | 1 | 20 | 0 | 0 | 3 |
| Banf1<sub>Thr2</sub>_Lys6 | 95 | 98 | 77 | 4 | 1 | 20 | 0 | 0 | 3 |
| Banf1<sub>Met1</sub>_Met1 | 0 | 0 | 0 | 0 | 0 | 0 | 0 | 0 | 0 |
| Banf1<sub>Met1</sub>_Thr2 | 0 | 8 | 10 | 1 | 1 | 19 | 0 | 0 | 0 |
| Banf1<sub>Met1</sub>_Res3 | 0 | 86 | 46 | 1 | 12 | 35 | 42 | 2 | 18 |
| Banf1<sub>Met1</sub>_Res4 | 0 | 88 | 52 | 0 | 11 | 34 | 2 | 0 | 10 |
| Banf1<sub>Met1</sub>_Gln5 | 96 | 99 | 76 | 4 | 1 | 22 | 0 | 0 | 2 |
| Banf1<sub>Met1</sub>_Lys6 | 96 | 99 | 78 | 4 | 1 | 20 | 0 | 0 | 2 |

Although Banf1 functions as a dimer, some human Banf1 variants are known or predicted to disturb the dimerization stability of Banf1 monomers [13]. Therefore, we explored if the phosphorylation impact on the N-terminal conformation of Banf1 is affected by its dimerization state, by performing MD simulations on unphosphorylated, mono-phosphorylated and di-phosphorylated Banf1_Met1_ and Banf1_Thr2_ monomer. Figure 8 displays the secondary structure variations with respect to time of the N-terminal residues in Banf1_Thr2_, which were analysed from one of the three MD simulation replicas of the corresponding Banf1_Thr2_ monomer. The corresponding results from the other two simulation replicas of Banf1_Thr2_ and the three simulation replicas of Banf1_Met1_ monomer are shown in Supplementary Figure S4. Here, we found that the secondary structure changes in the N-terminal region of a Banf1 monomer are overall consistent among all the three copies of simulations, suggesting the reliability of the MD results. Notably, due to the salt bridge interaction between pThr3 and Lys72, the N-terminal conformation of pSer4 Banf1_Met1_ corresponding to the secondary structure map within the magenta dashed line rectangular in Figure S4 differs from the N-terminal conformation of the representative structure of pSer4 Banf1_Met1_ (cyan) in Figure S3b1, but can also inhibits the DNA-binding capability of Banf1 by inducing steric clashes between Banf1 and DNA (Figure S5). The N-terminal domain of pSer4 Banf1_Met1_ overcomes this conformation to reform a helical structure after ∼60 ns and this conformation was not very stable in the other simulation replicas of Banf1. In this conformation (Figure S5), the phosphate group of pSer4 forms a salt bridge with the ε-amino group of Lys72, which agrees with the results by Marcelot et al. that the conformational restriction of the N-terminal domain of phosphorylated Banf1 is regulated by the dynamic network of salt bridge interactions involving pSer4 and pThr3 with positively charged residues located in nearby α1 helix and α6 helix [67]. In addition, the dynamics of the secondary structures of the N-terminal residues in pSer4 Banf1 and pBanf1 summarised from simulations of Banf1 monomers (Figure 8 and S4) agree with those analysed from simulations of Banf1 dimers (Figure 7 and S3). These results reveal that the phosphorylation impact on the N-terminal conformation of Banf1 is independent of Banf1’s dimerization state and stability, due to the fact that the N-terminal region is located on the opposite side of the dimerization surface in Banf1 (Figure 1A). This is further supported by the fact that phosphorylation of Banf1 does not affect its binding to emerin, which binds to Banf1 dimer via a site around its dimerization surface [17].

**Figure 8.**
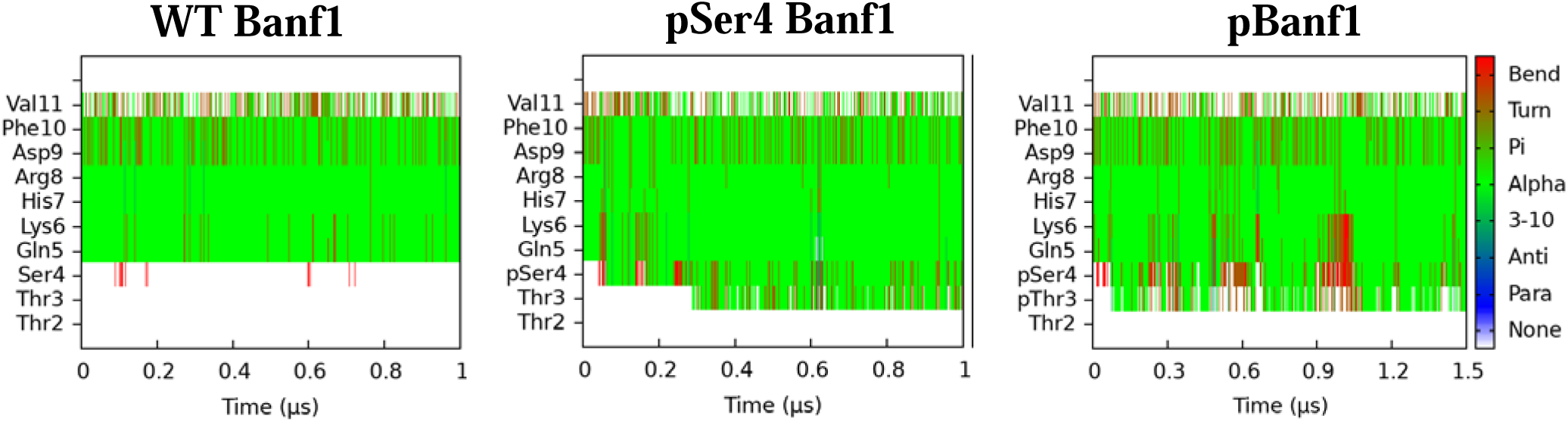
Secondary structure maps with respect to time of the N-terminal residues of Banf1_Thr2_, analysed from simulations of Banf1_Thr2_ monomer. The three secondary structure maps from left to right are for residues 2-11 of unphosphorylated, mono-phosphorylated and di-phosphorylated Banf1_Thr2_, respectively.

#### Phosphorylation alters hydrogen bonding interactions and induces new salt bridge interactions in Banf1

To explore the atomic-level mechanisms underlying the aforementioned findings that mono- and di-phosphorylation induce distinct secondary structure changes in the N-terminal region of Banf1, we performed hydrogen bonding and salt bridge analysis for the simulation trajectories of unphosphorylated, mono-phosphorylated and di-phosphorylated Banf1_Met1_ and Banf1_Thr2_ dimer. Table 5 displays some of the hydrogen bonds and salt bridges in the 6 systems that are related to changes in the N-terminal secondary structure and DNA binding surface of Banf1 induced by phosphorylation.

**Table 5.** Occupancies of some hydrogen bonds and salt bridges that are related to changes in the N-terminal conformation and DNA binding surface of Banf1 caused by phosphorylation. The information for salt bridges is highlighted in bold. The cut-off distance of salt bridges is 6 Å. The occupancies were averaged from two monomers of the dimer. OxP represents phosphate oxygens of the corresponding phosphate group.

| Acceptor | DonorH | Donor | WT<br>Banf1<sub>Thr2</sub> | pSer4<br>Banf1<sub>Thr2</sub> | pBanf1<sub>Thr2</sub> | WT<br>Banf1<sub>Met1</sub> | pSer4<br>Banf1<sub>Met1</sub> | pBanf1<sub>Met1</sub> |
| --- | --- | --- | --- | --- | --- | --- | --- | --- |
| (p)Ser4@O | Arg8@H | Arg8@N | 99 | 94 | 86 | 99 | 97 | 81 |
| Gln5@O | Asp9@H | Asp9@N | 94 | 94 | 73 | 94 | 95 | 70 |
| Lys6@O | Phe10@H | Phe10@N | 75 | 67 | 70 | 77 | 67 | 68 |
| Gly21@O | Lys6@HZ1/2/3 | Lys6@NZ | 79 | 75 | 78 | 79 | 76 | 74 |
| Leu23@O | Lys6@HZ1/2/3 | Lys6@NZ | 28 | 26 | 30 | 26 | 27 | 26 |
| Ile26@O | Lys6@HZ1/2/3 | Lys6@NZ | 31 | 28 | 30 | 28 | 28 | 39 |
| Ser4@OG | His7@H | His7@N | 90 |  |  | 92 |  |  |
| Thr3@O | Arg8@HE | Arg8@NE | 31 |  |  | 34 |  |  |
| Thr3@O | Arg8@HH12/21 | Arg8@NH1/2 | 33 |  |  | 40 |  |  |
| Thr3@OG1 | Arg8@HE | Arg8@NE | 32 |  |  | 28 |  |  |
| Thr3@OG1 | Arg8@HH12/21 | Arg8@NH1/2 | 28 |  |  | 28 |  |  |
| **pSer4@OxP** |  | **Arg8@NH1/NH2** |  | **98** | **78** |  | **95** | **73** |
| **pThr3@OxP** |  | **Lys72@NZ** |  |  | **57** |  |  | **51** |
| Thr2@O | Lys6@H | Lys6@N |  | 62 | 39 |  | 95 | 47 |
| (p)Thr3@O | His7@H | His7@N |  | 73 | 46 |  | 88 | 44 |
| Thr2@OG1 | Gln5@H | Gln5@N |  | 91 | 29 |  | 84 | 26 |
| pSer4@OxP | Thr2@HG1 | Thr2@OG1 |  | 94 | 39 |  | 85 | 16 |
| pThr3@O | Lys6@H | Lys6@N |  |  | 27 |  |  | 21 |
| pThr3@OxP | pThr3@H | pThr3@N |  |  | 37 |  |  | 47 |
| Met1@O | pSer4@H | pSer4@N |  |  |  |  | 7 | 26 |
| Met1@O | Gln5@H | Gln5@N |  |  |  |  | 6 | 20 |

In general, an α-helix is stabilised by hydrogen bonds between every backbone carbonyl group (C=O) and the backbone amide group (N-H) of amino acids located 4 residues later along the protein sequence, as exemplified by the N-terminal α1 helix of Banf1 in Figure 1B and 1C. As shown in Table 5, the occupancies of backbone hydrogen bonds (residue 4 with Arg8, and Gln5 with Asp9) that stabilise the N-terminal α1 helix are high (over 94%) in WT Banf1 and pSer4 Banf1. Further phosphorylation at Thr3 of pSer4 Banf1 significantly reduces the hydrogen bonding frequency by ∼20% between pSer4 and Arg8 and by ∼30% between Gln5 and Asp9, due to the alterations in electrostatic interactions between (p)Thr3 and nearby residues induced by phosphorylation of Thr3. (Table 5). This results in the significantly shortened N-terminal α1 helix observed in X-ray diffraction experiments [17] and highly frequently adopted by pBanf1_Met1_ and pBanf1_Thr2_ in MD simulations (Table 4, and Figure 7c2 and S3c2). Since Thr3 of Banf1 is relatively far away from residues Lys6 and Phe10, its phosphorylation does not affect hydrogen bonding interactions between the carbonyl group of Lys6 and the amide group of Phe10, which explains why both mono- and di-phosphorylation only induce local conformational changes in the vicinity of phosphorylation site(s) of Banf1. Similarly, since both mono- and di-phosphorylation only induce local conformational changes in the vicinity of phosphorylation site(s) of Banf1, they do not affect the restriction of the pocket, formed by the carbonyl groups of Gly21, Leu23 and Ile26, on the ε-amino group of Lys6; the ε-amino group of Lys6 hydrogen bonds to the carbonyl groups of Gly21, Leu23 and Ile26 with similar frequencies in WT Banf1, pSer4 Banf1 and pBanf1 (Table 5).

The N-terminal secondary structure of Banf1 is mainly determined by electrostatic interactions. In WT Banf1, the N-terminal conformation is confined by hydrogen bonds formed at two sides of its backbone (Table 5, Figure 7a1 and Figure S3a2): on the side close to the α6 helix (blue in Figure 1), there is a highly frequent hydrogen bond (90% for WT Banf1_Thr2_ and 92% for WT Banf1_Met1_) between the hydroxyl group of Ser4 and the backbone amide hydrogen of His7; on the other side, the guanidine group of Arg8 hydrogen bonds to the carbonyl oxygen and/or the hydroxyl group of Thr3. As shown in Table 5, these hydrogen bonds do not present in pSer4 Banf1 and pBanf1. Instead, due to the addition of the negatively charged phosphate group to Ser4 (and Thr3), mono- and di-phosphorylation induce some new hydrogen bonds and salt bridges in Banf1 (Table 5 and Figure 7 and S3), thus causing the conformational changes in its N-terminal domain. Particularly, the negatively changed phosphate oxygens of pSer4 form a highly frequent salt bridge with the positively charged guanidine nitrogens of Arg8 with a fraction of 98% and 95% in the simulations of pSer4 Banf1_Thr2_ and pSer4 Banf1_Met1_, respectively. These salt bridges occur relatively less frequent in pBanf1, with a propensity of 78% for pBanf1_Thr2_ and 73% for pBanf1_Met1_. An additional frequent salt bridge between the phosphate oxygens of pThr3 and the ε-amino nitrogen of Lys72 is observed in the simulations of pBanf1 (Table 5 and Figure 7 and S3), with a fraction of 57% and 51% for pBanf1_Thr2_ and pBanf1_Met1_, respectively. This salt bridge interaction presents in all the three representative conformations of pBanf1 (Figure 7 and S3) and was recently characterised by X-ray diffraction experiments for pBanf1_Thr2_ [17], demonstrating the reliability of our MD results.

Apart from inducing new salt bridge(s), mono- and di-phosphorylation of Banf1 result in additional backbone hydrogen bonds that stabilise an alpha helical structure. Specifically, with the help of the strong electrostatic interactions between the phosphorylated residues and the charged residues Lys72 and Arg75 in the nearby α6 helix [15], the carbonyl group of Thr2 and (p)Thr3 frequently hydrogen bond to the amide group of Lys6 and His7, respectively, in mono- and di-phosphorylated Banf1 (Table 5). The occurrence of these two hydrogen bonds is more frequent (over 50%) in pSer4 Banf1 than in pBanf1, due to the salt bridge interaction in pBanf1 between negatively charged pThr3 and the positively charged residues in the nearby α6 helix [17]. These two new backbone hydrogen bonds formed between residues with 4 residues away from each other along the sequence, stabilise additional helical structures in residues 2-4 of pSer4 Banf1 and pBanf1. This results in the phosphorylated Banf1 conformations with elongated N-terminal α1 helix frequently observed in our MD simulations (Figure 7b1, 7c1, S3b1 and S3c1) and co-crystalised in 7NDY [17]. The much lower frequencies of the two backbone hydrogen bonds in pBanf1 than in pSer4 Banf1 explain why residues 3-4 in pBanf1 adopted alpha helical structure with significantly lower frequencies than in pSer4 Banf1 (Table1 and Figure 7 and S3). The new N-terminal conformations of phosphorylated Banf1 with an elongated alpha helical structure were further stabilised by the frequent hydrogen bonds of the hydroxyl group of Thr2 with the backbone amide group of Gln5 and the phosphate group of pSer4 (Table 5, Figure 7b1, 7c1 and S3b1). These two hydrogen bonds are also much more stable in pSer4 Banf1 than in pBanf1, due to the salt bridge interaction between the phosphate group of pThr3 and the ε-amino group of Lys72 in pBanf1 [17].

As shown in Table 5 and displayed in Figure 7 and S3, further phosphorylation at Thr3 of pSer4 Banf1 induces new hydrogen bonding interactions, due to the addition of the negatively charged phosphate group to Ser4 (and Thr3) and their interactions with the nearby charged residues. Specifically, the backbone carbonyl group of pThr3 hydrogen bonds to the backbone amide group of Lys6 with a propensity of 27% and 21% in pBanf1_Thr2_ and pBanf1_Met1_, respectively, which were not observed in WT Banf1 and pSer4 Banf1. Similar occupancies were observed for hydrogen bonding interactions of the carbonyl group of Met1 in pBanf1_Met1_ with the backbone amide groups of pSer4 (26%) and Gln5 (20%), as displayed in Table 5 and Figure S3c3. These two hydrogen bonds were much less frequently observed in the simulation of pSer4 pBanf1_Met1_. Interestingly, the phosphate oxygens of pThr3 form an intraresidue hydrogen bond with its backbone amide group with a frequency of 37% and 47% in pBanf1_Thr2_ and pBanf1_Met1_ (Table 5 and Figure S3c1) respectively.

In summary, due to the addition of the negatively charged phosphate group to Ser4 (and Thr3), both mono- and di-phosphorylation alter the atomic interactions around the phosphorylation site(s), by eliminating some hydrogen bonds present in WT Banf1 and inducing new electrostatic interactions, including hydrogen bonds and salt bridges. These alterations in interactions between residues near the phosphorylation site(s) result in significant local conformational changes in the N-terminal region of Banf1, which is independent of whether Banf1 has Met1. Particularly, all three representative conformations of pBanf1_Met1_ and pBanf1_Thr2_ are stabilised by two salt bridges (Figure 7 and S3), including i) a salt bridge between the phosphate oxygens of pSer4 and the guanidine nitrogens of Arg8; and ii) a salt bridge between the phosphate oxygens of pThr3 and the ε-amino nitrogen of Lys72 (characterised for pBanf1_Thr2_ in Ref. [17]). In contrast, the N-terminal conformation of pSer4 Banf1_Met1_ and pSer4 Banf1_Thr2_ are unique and are stabilised by a salt bridge between the phosphate oxygens of pSer4 and the guanidine nitrogens of Arg8, and by 4 hydrogen bonds (Table 5 and Figure 7b1 and S3b1), including i) a backbone hydrogen bond between the carbonyl group of Thr2 and the amide group of Lys6; ii) a backbone hydrogen bond between the carbonyl group of Thr3 and the amide group of His7; iii) a hydrogen bond between the hydroxyl hydrogen of Thr2 and the phosphate group of pSer4; and iv) a hydrogen bond between the hydroxyl oxygen of Thr2 and the amide group of Gln5.

#### Phosphorylation changes the morphology and electrostatic potential distribution of the DNA binding surface of Banf1

Binding of a protein to DNA is closely regulated by the morphology and chemical properties of its DNA binding surface. To explore how phosphorylation of Banf1 destroys its DNA-binding capability, we investigated alterations in the morphology and electrostatic potential distribution of the DNA binding surface of Banf1 caused by its mono- and di-phosphorylation. Figure 9 and Supplementary Figure S6 display the hydrophobicity surface representation of the representative conformations of Banf1_Thr2_ and Banf1_Met1_ shown in Figure 7 and S3, respectively. As shown in Figure 1B and 6a, surface of the N-terminal residues 2-6 in WT Banf1_Thr2_ co-crystalised in 2BZF is approximately flat at the adopted perspective. Particularly, surfaces of Ser4, Gln5 and Lys6 (highlighted in yellow) in WT Banf1 are connected to each other (Figure 9a1) and their backbones are simultaneously directly exposed to the solvent accessible for DNA backbone phosphates (Figure 1), thus allowing WT Banf1 to bind to DNA double helix strongly. Specifically, the backbone amide group of Gln5 and Lys6 form close interactions with the backbone phosphate oxygens of the same DNA nucleotide (Figure 1C), with a hydrogen bond length of 1.8 Å and 2 Å, respectively [11]. In addition, the sidechain of Ser4 was characterised to form a water bridge with the backbone phosphate of the nucleotide adjacent to the nucleotide hydrogen bonding to Gln5 and Lys6 [11]. Figure 9a1 shows the hydrophobicity surface representation of the representative conformation (with an occupancy of 84%) of WT Banf1_Thr2_, which is close to that of WT Banf1_Thr2_ co-crystalised with DNA in 2BZF. This agrees with the fact that the DNA binding surface of WT Banf1 mainly consists of the backbone of some residues (e.g., Gln5, Lys6, Gly25, Ile26, Gly27, Leu30 and Ala71), the sidechain of some residues (Ser4, Lys6, Asn70, Lys72, Gln73 and Asp76) which are relatively stable in solution due to the intramolecular electrostatic interactions, and the sidechain of some hydrophobic residues (e.g., Ala24 and Leu 30) that are involved in intramolecular hydrophobic interactions. Notably, lengthy side chains of residue Arg85 that also contribute to WT Banf1’s DNA-binding surface are highly flexible in solution and hence are unlikely to always maintain exactly the same orientation as when Banf1 is bound to DNA. Upon mono- and di-phosphorylation, the morphologies of Banf1’s DNA binding surfaces are severely altered, as shown in Figure 9 and S6; surfaces of the N-terminal region involved in DNA binding are not flat in both mono- and di-phosphorylated Banf1. Unlike connected to each other in WT Banf1_Thr2_ (Figure 9a and 9a1), surfaces of Ser4 and Lys6 are far away from each other in all the representative conformations of pSer4 Banf1 and pBanf1. Clearly, the simultaneous accessibility to DNA of Ser4, Gln5 and Lys6 are blocked by Thr2 (white) in pSer4 Banf1_Thr2_ and by Thr2 and pThr3 (red) in pBanf1_Thr2_ (Figure 9), and by Met1 (pink), Thr2 and (p)Thr3 in pSer4 Banf1_Met1_ and pBanf1_Met1_ (Figure S6). These significant changes in the morphology of Banf1’s DNA binding surface cause severe steric clashes in Banf1-DNA binding, thus preventing phosphorylated Banf1 from binding to DNA.

**Figure 9.**
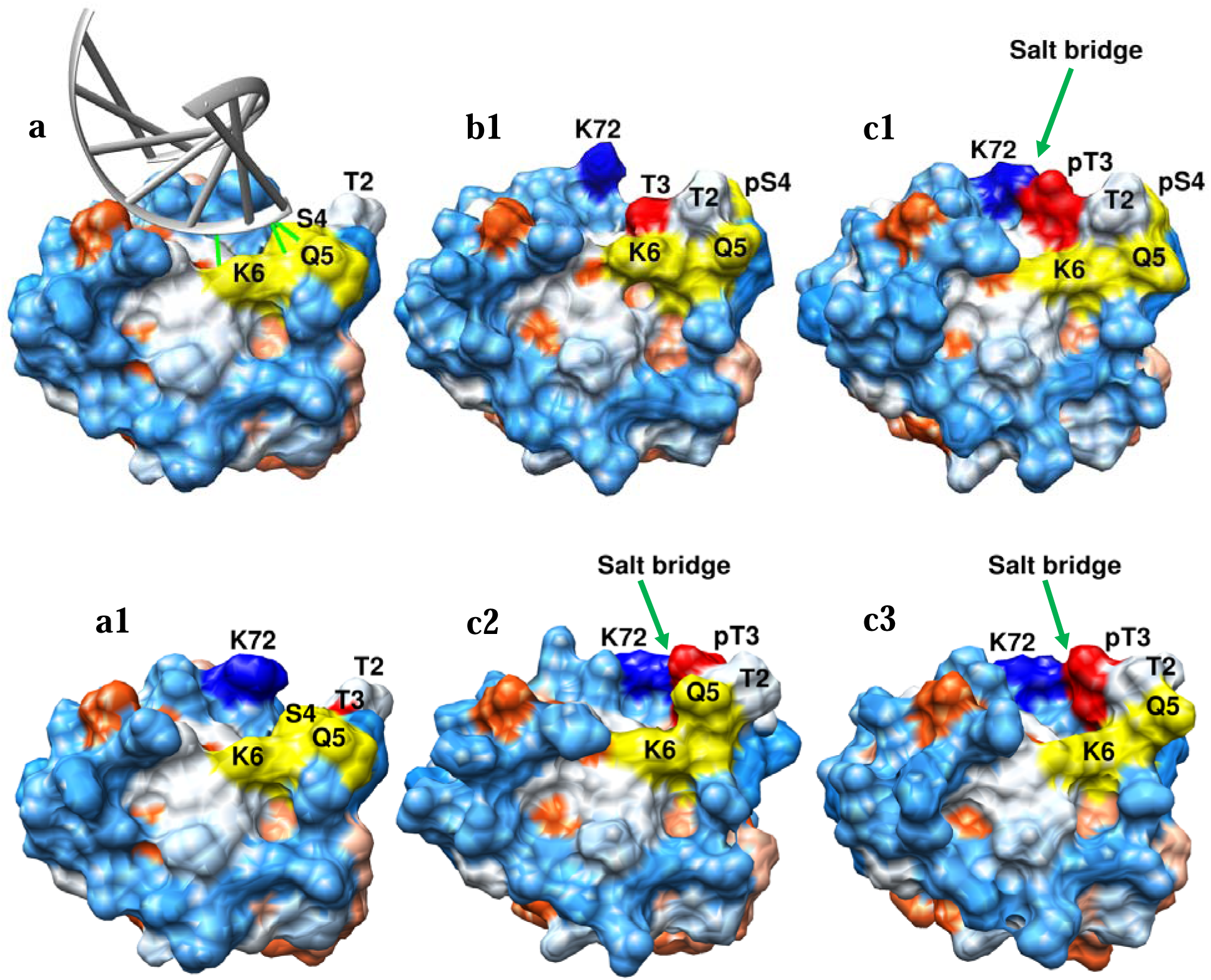
Hydrophobicity surface representation of the representative conformations of Banf1_Thr2_ displayed in Figure 7, where blue indicates the surface is hydrophilic and red is for hydrophobic surface. a is the hydrophobicity surface representation of Banf1_Thr2_ bound to DNA (coloured in dark gray) crystalised in 2BZF and displayed in Figure 1B. a1 and b1 are for the representative conformation of WT Banf1_Thr2_ and pSer4 Banf1_Thr2_, respectively. c1, c2, and c3 are for the three representative conformations of pBanf1_Thr2_. Figure numbers for each representative conformation are consistent with those in Figure 7. The perspective for each conformation are consistent as reflected by the location of K6 whose ε-amino group is restricted by a pocket formed by Gly21, Leu23 and Ile26 in the simulations of all systems. N-terminal residues 4-6 involved in DNA binding are highlighted in yellow. Residues Thr2, (p)Thr3 and Lys72 are coloured in white, red and blue, respectively. The locations of salt bridges are directed by green arrows. In c2 and c3, surfaces of pSer4 are behind the surfaces of Thr2 and Gln5, and are not visible in this perspective.

Figure 10 and Supplementary Figure S7 show the electrostatic potential surfaces of Banf1_Thr2_ and Banf1_Met1_ calculated based on Coulomb’s Law, respectively. Consistent with conclusions from Figure 9 and Supplementary Figure S6, the morphologies of Banf1’s DNA binding surfaces around the N-terminal region are severely altered by phosphorylation. As illustrated in Figure 10a1 and Figure S7a2, the DNA binding surfaces of WT Banf1_Thr2_ and WT Banf1_Met1_ are highly electropositive (blue), which enables the protein to bind strongly to the electronegative DNA phosphate backbone. Upon phosphorylation, the electrostatic potential distribution of Banf1’s DNA binding surfaces is remarkably changed, as shown in Figure 10 and S7. Particularly, the electrostatic potential surfaces of Banf1’s N-terminal region in both mono- and di-phosphorylated Banf1 are very different compared to WT Banf1, due to the distinct N-terminal conformational changes and the altered electrostatic interactions. Interestingly, the electrostatic potential of other regions comprising the DNA binding surface of Banf1 is either much less electropositive or almost neutral in pBanf1, due to the significant electrostatic potential modifications in the N-terminal region and the salt bridge between Lys72 and negatively charged pThr3. These changes in the electrostatic potential distribution of the DNA binding surface of Banf1 cause unfavourable interactions between Banf1 and DNA, thus preventing the occurrence of their binding. We note that, in this section, the perspective for all hydrophilic surface representations in Figure 9 and Figure S6 are the same, and the perspective for all electrostatic potential surface representations in Figure 10 and Figure S7 are consistent.

**Figure 10.**
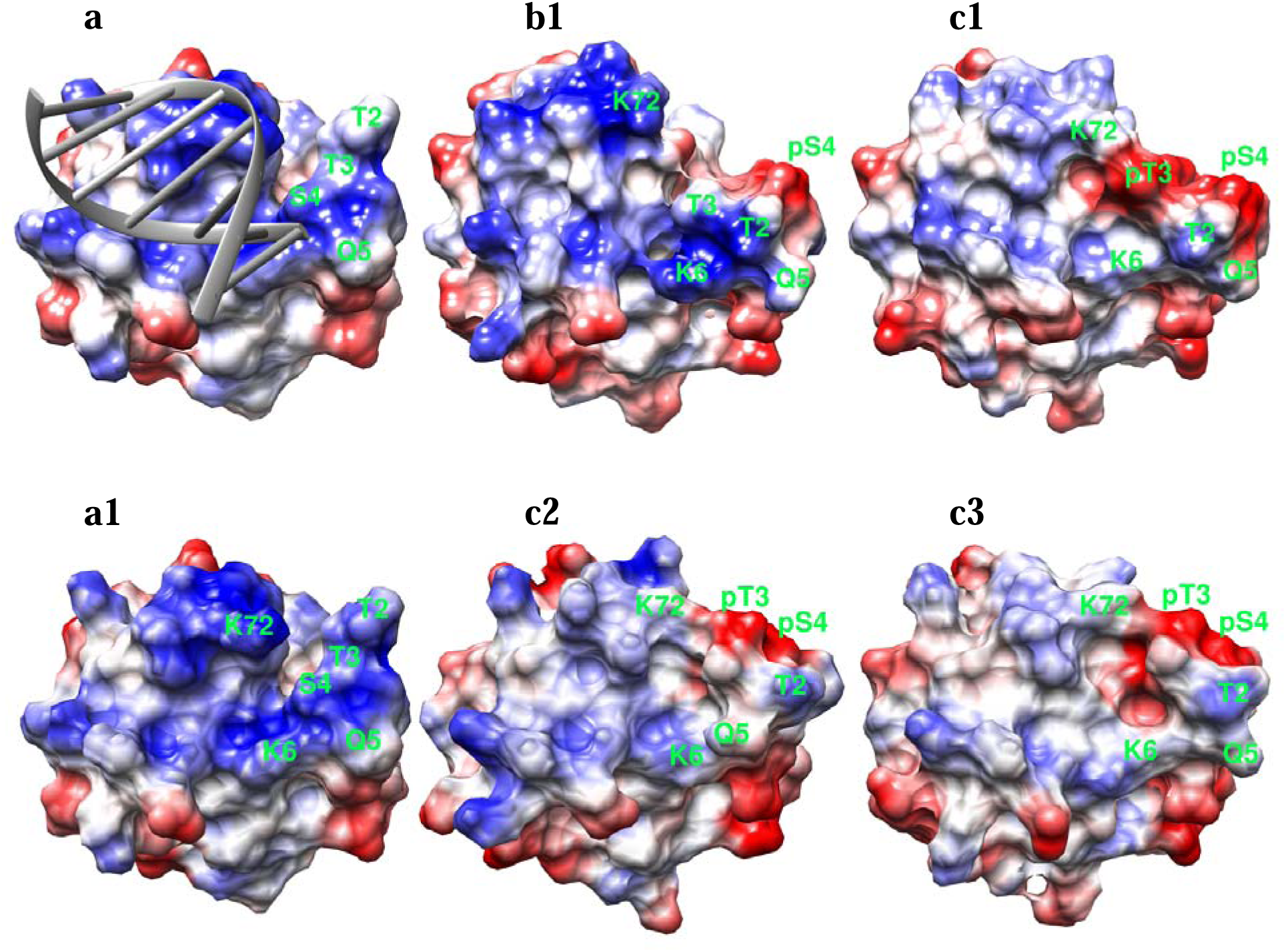
Electrostatic potential surface representation of the representative conformations of Banf1_Thr2_ displayed in Figure 7 and 9, where blue, white and red correspond to electropositive, electroneutral and electronegative surfaces, respectively. a is the electrostatic surface representation of Banf1_Thr2_ bound to DNA (coloured in dark gray) crystalised in 2BZF and displayed in Figure 1B. a1 and b1 are for the representative conformations of WT Banf1_Thr2_ and pSer4 Banf1_Thr2_, respectively. c1, c2, and c3 are for the three representative conformations of pBanf1_Thr2_. Figure numbers for each representative conformation are consistent with those in Figure 7 and 9.

### Binding of phosphorylated Banf1 with DNA

To study mono- and di-phosphorylation effect on Banf1’s DNA binding capability, we docked DNA from 2BZF to the representative structures of pSer4 Banf1_Thr2_ and pBanf1_Thr2_ displayed in Figure 7. Here we did not consider the binding of DNA to Banf1_Met1_, given that i) Met1 of Banf1 is not involved in DNA binding, ii) Banf1 has Thr2 with a small sidechain, it is natural to consider that the majority of Met1 is processed by methionine aminopeptidase (MAP), and iii) mechanisms that define N-terminal phosphorylation in Banf1 impairs its DNA-binding capability are independent of whether Banf1 has Met1 cleaved.

Table 6 displays the best HADDOCK scores of DNA to unphosphorylated Banf1_Thr2_, pSer4 Banf1_Thr2_ and pBanf1_Thr2_, as well as the corresponding docked Banf1_Thr2_-DNA complex structures. As shown in Table 6, both mono- and di-phosphorylation of Banf1 remarkably reduce HADDOCK score of DNA to Banf1, due to the shifted binding poses compared to that co-crystalised in 2BZF and to the elimination of the electrostatic interactions (e.g., hydrogen bonds in green line) between the DNA phosphate backbone and the N-terminal residues Ser4, Gln5 and Lys6 of Banf1 resulted from the N-terminal secondary structure changes in phosphorylated Banf1.

**Table 6.**
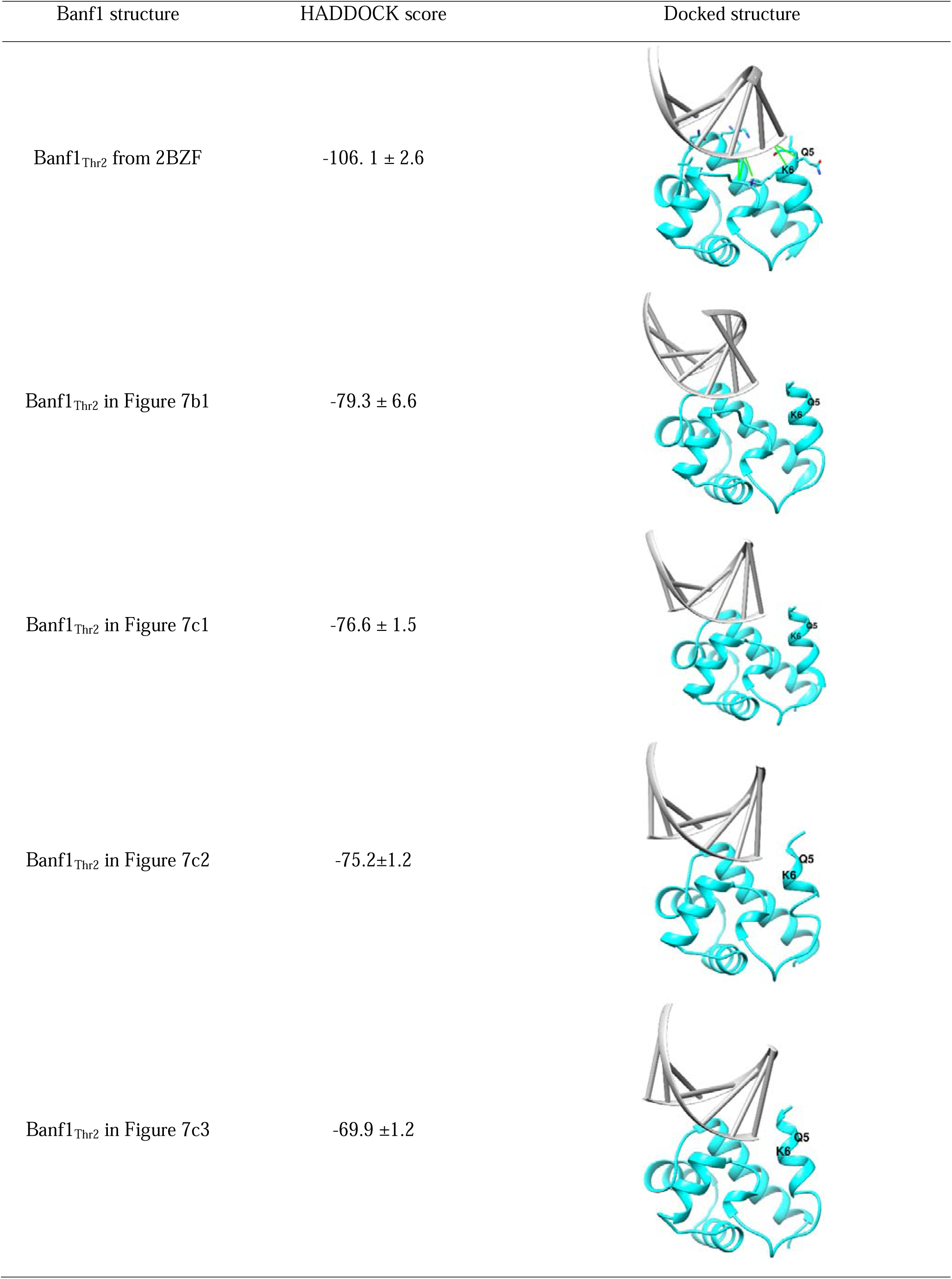
Docking scores from HADDOCK 2.4 of Banf1 and DNA and the corresponding docked complex structures. DNA is shown in gray ladder, and Banf1 structures in cyan ribbon.

| Banf1 structure | HADDOCK score | Docked structure |
| --- | --- | --- |
| Banf1<sub>Thr2</sub> from 2BZF | -106.1 ± 2.6 | |
| Banf1<sub>Thr2</sub> in Figure 7b1 | -79.3 ± 6.6 | |
| Banf1<sub>Thr2</sub> in Figure 7c1 | -76.6 ± 1.5 | |
| Banf1<sub>Thr2</sub> in Figure 7c2 | -75.2 ± 1.2 | |
| Banf1<sub>Thr2</sub> in Figure 7c3 | -69.9 ± 1.2 | |

To further explore the DNA-binding capability of phosphorylated Banf1, we performed MD simulations to check the stability of the top-scoring docked Banf1-DNA complexes displayed in Table 6. Our MD results showed that DNA molecule shifted from its initial docked poses in the Banf1-DNA complexes very quickly (within 20 ns) during the MD simulations, suggesting that these docked Banf1-DNA poses are not stable in solution. This indirectly suggests that the DNA-binding capability of pSer4 Banf1 and pBanf1 with their representative N-terminal structures are severely impaired. This result agrees with the experimental findings that pBanf1 binds to DNA with a 5000-folds lower affinity [17]. Notably, the detaching route and time may be related to the applied force fields for the protein, the phosphorylated residues and the DNA molecules [69]. For example, OL15 force field and the RESP partial charge method were found to be more stable and reliable for modelling the G-quadruplex systems. Here, we used the ff99SB force field for proteins and DNA, as the force fields for pSer4 and pThr3 were derived in consistent with the ff99SB force field.

## Discussion

Marcelot and co-workers revealed that di-phosphorylation of Banf1 destroys its DNA-binding capability by inducing steric clashes due to critical conformational changes of pBanf1’s N-terminal domain, which are confined by the electrostatic interactions between both phosphorylated residues (pThr3 and pSer4) and the charged residues Lys72 and Arg75 in the nearby α6 helix [17]. Importantly, there is a discrepancy in the length characterised in this study for the N-terminal α1 helix of pBanf1 [17], suggesting that the N-terminal secondary structure of in pBanf1 is relatively dynamic. Although pBanf1 was recently co-crystalised with emerin (PDB ID: 7NDY), the crystalised structure of pSer4 Banf1 is not yet available, possibly due to experimental difficulties. Here, we started with studying the detailed atomic bases of unphosphorylated Banf1-DNA binding, followed by exploring the impact of mono- and di-phosphorylation of Banf1 on these atomic bases. Specifically, we use the conformation of pBanf1 characterised by Marcelot et al. in 7NDY to validate the reliability of our modelling approaches and the force fields for the proteins, pThr3 and pSer4 to capture the N-terminal secondary of phosphorylated Banf1. By applying the validated MD approaches and force fields, we further disclosed the dynamics of the N-terminal secondary structure of pBanf1 and its DNA-binding surface, and revealed the molecular- and atomic-level mechanisms underpinning the phenomenon that mono-phosphorylated Banf1 with a N-terminal helical conformation is not capable of binding to DNA.

### Detailed atomic bases of the binding between unphosphorylated Banf1 and DNA

Met1 in Banf1 does not directly interact with DNA and whether Banf1 has Met1 at its N-terminus does not affect its DNA-binding activity. Here, we identified some new transient hydrogen bonds and new water-mediated hydrogen bonds between Banf1 and DNA that have not been reported in the literature. These direct and indirect hydrogen bonds were not characterised by X-ray crystallography, possibly because they are much less stable than the experimentally characterised 9 hydrogen bonds as reflected by their much lower occupancies.

### The molecular- and atomic-level mechanisms by which mono-phosphorylation and di-phosphorylation of Banf1 inhibit its binding to DNA

MD simulations confirmed our speculation that the N-terminal secondary structure of pBanf1 is relatively dynamic but is different from that of WT Banf1, which prevents pBanf1 from binding to DNA. The relative dynamic nature of the pBanf1 N-terminal domain might facilitate the dephosphorylation event of pBanf1 which is essential for the reassembly of nuclear envelope. At the molecular level, our modelling results agree with the X-ray diffraction results [17] in that di-phosphorylation of Banf1 inhibits its binding to DNA by inducing steric clashes due to the N-terminal conformational changes (https://www. youtu.be/Mn2o20m-J3g) [17]. Our results extend the experimental results in Ref [17] by uncovering the unfavourable electrostatic potential of the DNA binding surface in pBanf1. At the atomic level, our results align with the experimental results [17] in that the salt bridge between pThr3 in the N-terminal region and Lys72 in nearby α6 helix characterised for pBanf1_Thr2_ by X-ray (PDB ID: 7NDY) exists in all of the representative conformations of pBanf1 (Figure 7 and S3; Table 4). These results demonstrate the reliability of the force field applied for the protein, including the phosphorylated residues in our MD simulations to capture the dynamics of the N-terminal secondary structure of Banf1. Our results further extend the published experimental data [17] by revealing a new and strong salt bridge between the phosphate group of pSer4 and the guanidine group of Arg8 in pBanf1 (Figure 7 and S3). It is these two salt bridges located at two different sides of the N-terminal backbone of pBanf1 that restrict the N-terminal secondary structure of pBanf1, which in turn induces steric clashes and unfavourable interactions between Banf1 and DNA.

Notably, NMR experiments by Marcelot et al. found a significant reduction of the conformational dynamics in the N-terminal region of Banf1 upon its phosphorylation [17, 67]. However, no significant conformational change induced by phosphorylation was detected in the crystal state [17, 67]. This suggests that the crystal state may only partly reflects the conformational behaviour of the N-terminal domain of Banf1 [67]. Coincident with the submission of this paper, Marcelot et al. reported that the flexibility of the N-terminal domain of Banf1 in solution decreases with an increase in pH, and is gradually reduced by phosphorylation of Ser4 followed by phosphorylation of Thr3 [67]. Specifically, mono-phosphorylation of Banf1 confines the space for the conformational ensembles of Banf1, and this restriction is more significant when pThr3 is also phosphorylated [67]. In this paper, we focused on the detailed atomic bases of Banf1-DNA binding, and the impacts of mono-phosphorylation of Banf1 on its N-terminal secondary structure as well as its DNA binding surface, hoping to provide insights into why mono-phosphorylated Banf1 barely binds DNA. For comparison, we did clustering analysis and superimposed the top 100 representative conformations (in ribbon) of unphosphorylated (Figure S8A), mono-phosphorylated (Figure S8B) and di-phosphorylated (Figure S8C) Banf1_Thr2._ Our results qualitatively agree with the findings by Marcelot et al. that phosphorylation confines the ensemble space that can be occupied by Banf1’s N-terminal region [67].

The salt bridge between the phosphate oxygens of pSer4 and the guanidine nitrogens of Arg8 is even stronger in pSer4 Banf1 than that in pBanf1, which is almost rigid in pSer4 pBanf1_Thr2_ and pSer4 pBanf1_Met1_ (Table 5). Unlike pBanf1, where Thr3 is phosphorylated into pThr3, whose phosphate group can form a salt bridge with the nearby Lys72, Thr3 in pSer4 Banf1 is neutral and has a shorter sidechain compared to pThr3, thus not allowing it to form strong electrostatic interactions with the ε-amino group of Lys72 located in α6 helix. Instead, its adjacent residue Thr2 in pSer4 Banf1 forms hydrogen bonding interactions with the backbone amide group of Gln5 and the phosphate oxygens of pSer4 via its hydroxyl group (Figure 7b1 and S3b1). It is these two strong hydrogen bonds and the almost solid salt bridge between pSer4 and Arg8 that lock the conformation of the N-terminal region of pSer4 Banf1 as a helical structure, which alters the DNA binding surface of pSer4 Banf1, induces steric clashes and unfavourable interactions between pSer4 Banf1 and DNA and consequently prevents it from binding to DNA. For the first time, this study reveals the atomic-level mechanisms underlying the experimental observations that mono-phosphorylation at Ser4 of Banf1 significantly inhibits its binding to DNA [17, 18]. It is worth noting that increasing the negative charge of Banf1 N-terminal region progressively shifts Banf1 N-terminal conformations towards a more rigid and unique structure [17, 67] (Figure S8), although the N-terminal structure of pSer4 Banf1 seems relative more stable than that of pBanf1 when they both adopt a helical conformation (Table 1, Figure 7 and S3). Collectively, mono-phosphorylation of Banf1 destroys its DNA-binding capability via the same mechanism as di-phosphorylation does at the molecular level, i.e., via steric clashes and unfavourable interactions between the protein and DNA originated from critical conformational changes in Banf1 N-terminal region (Figure 11). This agrees with the experimental findings that pSer4 Banf1 phosphomimetics are not able to bind DNA molecules [17, 18]. However, the atomic-level interactions that are responsible for the N-terminal secondary structure changes in pSer4 Banf1 and pBanf1 are distinctly different (Table 5 and Figure 7 and S3).

**Figure 11.**
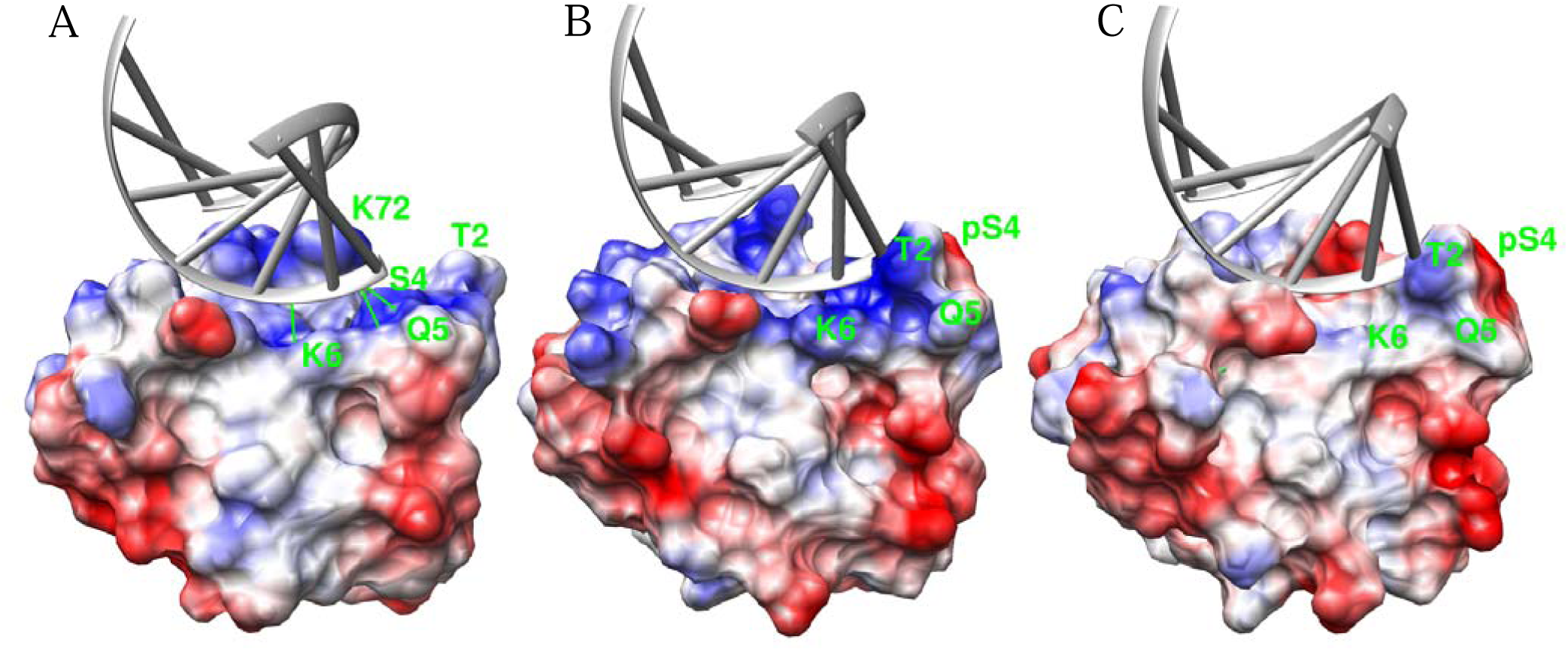
Mechanisms of how mono- and di-phosphorylation of Banf1 inhibit its binding to DNA. A is electrostatic potential surface representation of Banf1_Thr2_ bound to DNA which is co-crystalised in 2BZF and displayed in Figure 1B. B is the electrostatic potential surface representation of the representative conformation of mono-phosphorylated Banf1_Thr2_ in Figure 7b1 and 9b1 with the DNA in 2BZF after superimposing Banf1 in the two structures. C is the electrostatic potential surface representation of a representative conformation of di-phosphorylated Banf1_Thr2_ in Figure 7c1 and 9c1 with the DNA in 2BZF after superimposing Banf1 in the two structures. Hydrogen bonds between DNA backbone and Banf1 N-terminal residues are displayed in green solid lines in A. DNA is coloured in dark gray and residue numbers in green. Perspective of this figure is the same as that of Figure 9. Clearly, both mono- and di-phosphorylation alter the DNA binding surface of Banf1, which indues steric clashes and unfavourable interactions between DNA and Banf1, thus preventing their binding.

### Phosphorylation abrogates Banf1-DNA binding in both with and without Met1

Interestingly, Banf1_Met1_ [70] and Banf1_Thr2_ [71] were both detected by mass spectrometry in Jurkat cells [72] and Hela cells [73], respectively. Here, we revealed that mono- and di-phosphorylation inducing changes in the conformation and DNA binding surface of Banf1 are independent of whether it contains Met1 on the N-terminus, suggesting that phosphorylation of both forms of Banf1 prevents it from binding to DNA. This demonstrates that phosphorylation and dephosphorylation regulate the binding of DNA to Banf1_Met1_ and Banf1_Thr2_ in a similar way; unphosphorylated Banf1_Met1_ and Banf1_Thr2_ bind to DNA, whereas mono- and di-phosphorylation prevent the binding between DNA and Banf1_Met1_ and Banf1_Thr2_. This is consistent with the fact that both forms of Banf1 exist and function well in cells.

### Limitations and future work

Here, we applied classic MD simulations to characterise the N-terminal secondary structure and the DNA-binding surface of Banf1 with three different phosphorylation statuses, because: i) within the simulation length of 1/1.5 <u>μs adopted</u> in this manuscript, classic MD simulations are capable of capturing the secondary structure changes of Banf1’s N-terminal region induced by both mono- and di-phosphorylation; ii) the secondary structure changes of Banf1’s N-terminal domain with a certain phosphorylation status are qualitatively consistent in the simulation of Banf1 dimer and the three simulation replicas of Banf1 monomer; iii) the exact percentage of the representative conformations of Banf1 are not the focus of this manuscript. Enhanced sampling technics such as metadynamics and Gaussian accelerated MD simulations can provide a larger sampling space than the classic MD simulations used in this manuscript. For example, metadynamics can be used to improve the sampling space of the N-terminal conformation of Banf1, and to determine the free energies required for overcoming the representative conformations of Banf1.

Unlike WT Banf1-DNA complex which is stable with the binding pose co-crystalised in 2BZF in multiple 1-μs-long simulations with different atomic velocities, both mono- and di-phosphorylated Banf1 bind to DNA with a significantly lower docking score and the docked pSer4 Banf1-DNA and pBanf1-DNA complexes are not stable in solution in our MD simulations. Since Marcelot et al. [17] found that pBanf1 binds DNA with a Kd value of 11 ± 2 μM [17], pBanf1 should in principle binds DNA in solution, although with a significantly lower binding affinity than unphosphorylated Banf1. The weak binding of pBanf1 to DNA might be conferred by the flexibility of pBanf1’s N-terminal domain [17, 67], where the phosphorylated Banf1 with a severe N-terminal secondary structure change (e.g., from random loop to helix) cannot bind to DNA whereas some of the phosphorylated Banf1 with a similar N-terminal geometry as the N-terminal structure of unphosphorylated Banf1 are capable of binding to DNA to some extent. The ideal way to simulate if/how phosphorylated Banf1 binds to DNA is to randomly put phosphorylated Banf1 and DNA far away from each other in solution and check if/how they bind. In this scenario, DNA theoretically has chances to meet any of the phosphorylated Banf1’s N-terminal conformation. This scenario is out of the reach of classic MD simulations, due to the large size of the DNA molecule and the high flexibility of Banf1’s N-terminal domain. This study only focuses on exploring why already phosphorylated Banf1 barely binds to DNA in the nucleus and thus locates in the cytoplasm. Future work is required to uncover the unbinding mechanisms from DNA of Banf1 upon its mono- and di-phosphorylation, by simulating the induced fit of pSer4 Banf1-DNA and pBanf1-DNA complexes with a starting structure where Ser4 (and Thr3) is mutated in the Banf1-DNA complex co-crystalised in 2BZF.

## Conclusions

In this paper, we performed systematic and comprehensive molecular docking and molecular dynamics simulations, to explore i) detailed atomic bases responsible for the unphosphorylated Banf1-DNA binding, and ii) the dynamics of the N-terminal secondary structure and the DNA binding surface (morphology and electrostatic potential) of un-phosphorylated, mono-phosphorylated and di-phosphorylated Banf1. Our study identified new intermolecular hydrogen-bonding and water-bridge interactions in the Banf1-DNA complex. Moreover, our results demonstrated that the N-terminal secondary structures of pSer4 and pBanf1 are relatively dynamic. Additionally, this study unravelled how mono-phosphorylation at Ser4 induces distinct conformational changes in the vicinity of the phosphorylation site of Banf1, which results in significant changes in the morphology and electrostatic potential of its DNA binding surface. These critical alterations in the DNA binding surface of Banf1 eliminate its DNA-binding capability. This molecular-level mechanism of destroying the DNA-binding capability of Banf1 is the same for pSer4 Banf1 and pBanf1 and is independent of Banf1’s dimerization state and whether Banf1 has Met1 cleaved on the N-terminus. However, at the atomic level, we revealed that the relatively dynamic N-terminal secondary structure of pBanf1 is confined by two salt bridges at both sides of its backbone, whereas the pSer4 Banf1 N-terminal secondary structure is confined to a unique conformation by a salt bridge and a number of hydrogen bonds.

## Author Contributions

M.T. designed the project, did the modelling and wrote the manuscript. D.J.R. provided supervision. D.J.R. and E.B. provided resources. E.B., D.J.R., R.J.W., A.S., X.Q.N. Z.L.L., J.W.W. and K.J.O. provided precious discussions, comments and revisions. All authors contributed to the work and approved the publication of the manuscript.

## Conflict of Interest

The authors declare competing financial interests; E.B., D.J.R. and K.J.O. are founders of Carpe Vitae Pharmaceuticals. E.B., K.J.O. and D.J.R. are inventors on provisional patent applications filed by Queensland University of Technology.

## Acknowledgement

D.J.R acknowledges the financial support from William and Hilde Chenhall Research Trust. The High Performance Computing resources provided by Queensland University of Technology (QUT) are gratefully acknowledged. K.J.O acknowledges the Queensland Health Senior Clinical Research Fellowship (K.J.O) and a Yancoal research grant. X.Q.N. acknowledges the National Natural Science Foundation of China (81960741, 82160770), Outstanding Young Scientific and Technological Talents Project of Guizhou Province (2021-5639), and scholarships from the China Scholarship Council (CSC-202008520012).

---

**Figure S1.**
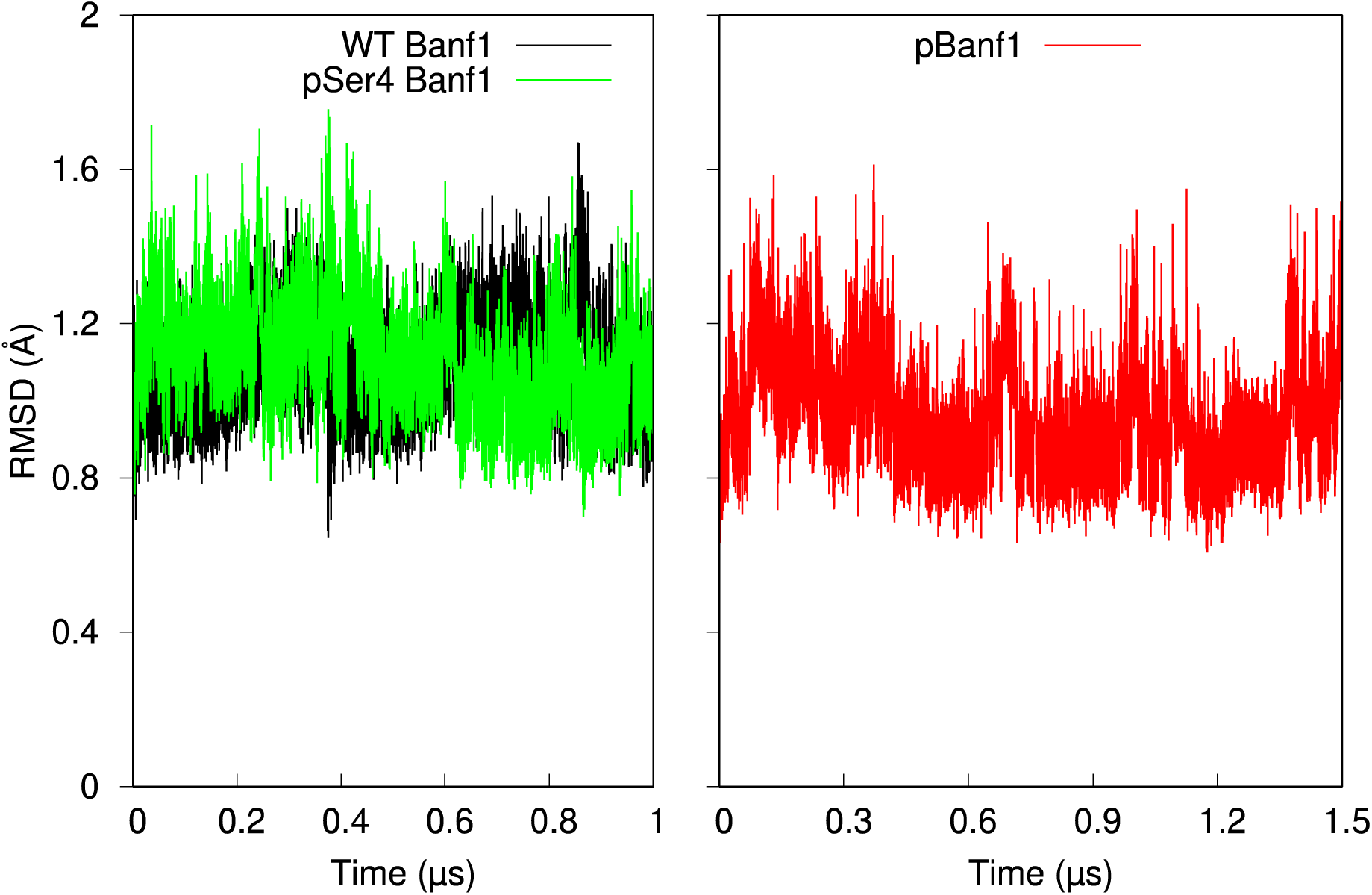
RMSD variations with respect to time of residues 7-89 in Banf1_Met1_, where WT Banf1 is for unphosphorylated Banf1_Met1_, pSer4 Banf1 for mono-phosphorylated Banf1_Met1_, and pBanf1 for di-phosphorylated Banf1_Met1._

**Figure S2.**
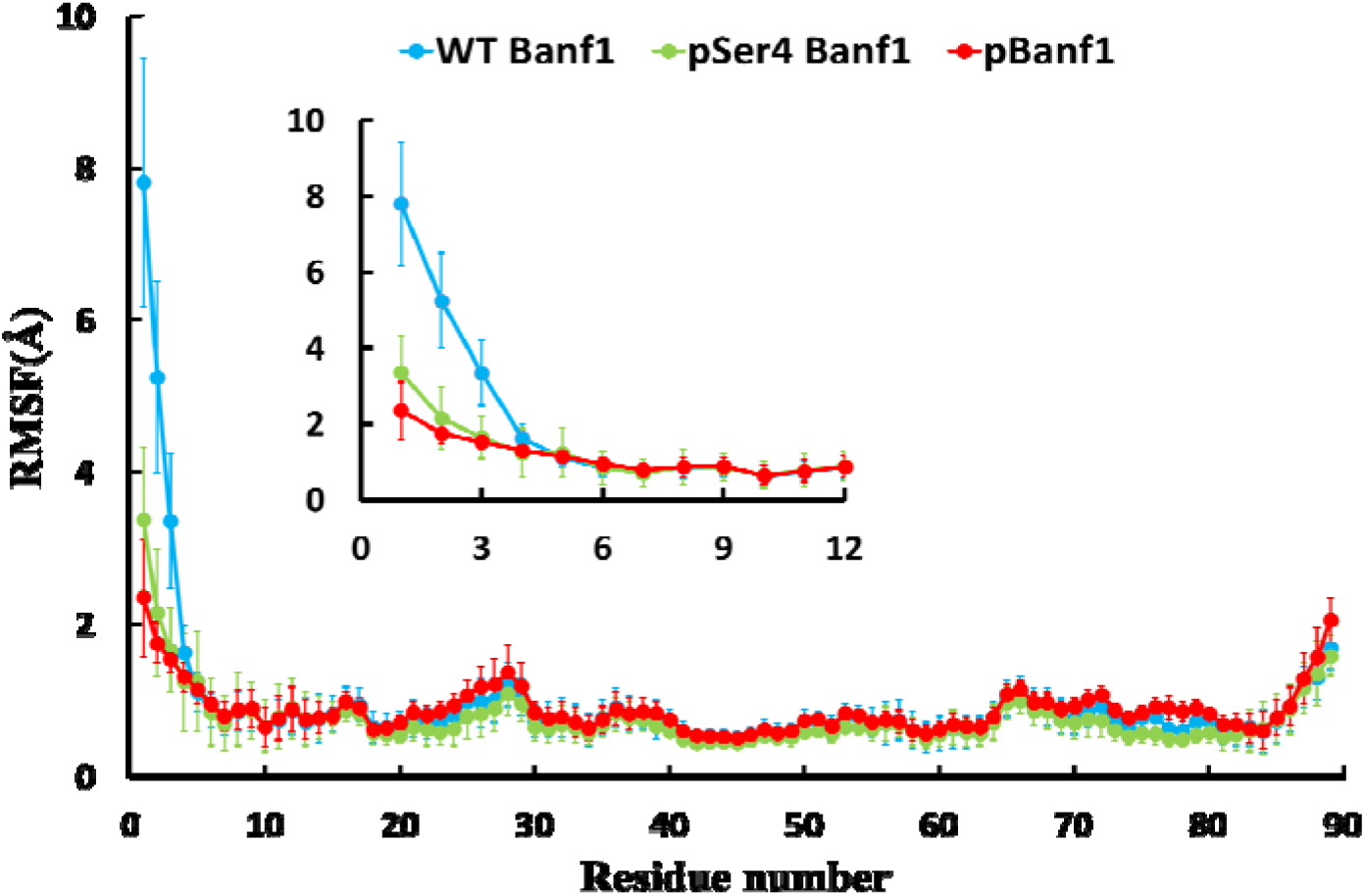
RMSF and its standard deviation of residues 1-89 in Banf1_Met1_ and its zoom in view incorporating the N-terminal residues 1-12 in Banf1_Met1_. The RMSF values were averaged from the RMSF of the two Banf1_Met1_ promoters in the simulation of Banf1_Met1_ dimer and the RMSF of the Banf1_Met1_ monomer in the three simulation replicas of Banf1_Met1_ monomer. Trajectories from the initial 100 ns simulations of unphosphorylated Banf1_Met1_ was discarded and the trajectories of simulations before the mono- and di-phosphorylated Banf1_Met1_ form an additional N-terminal helix were ignored.

**Figure S3.**
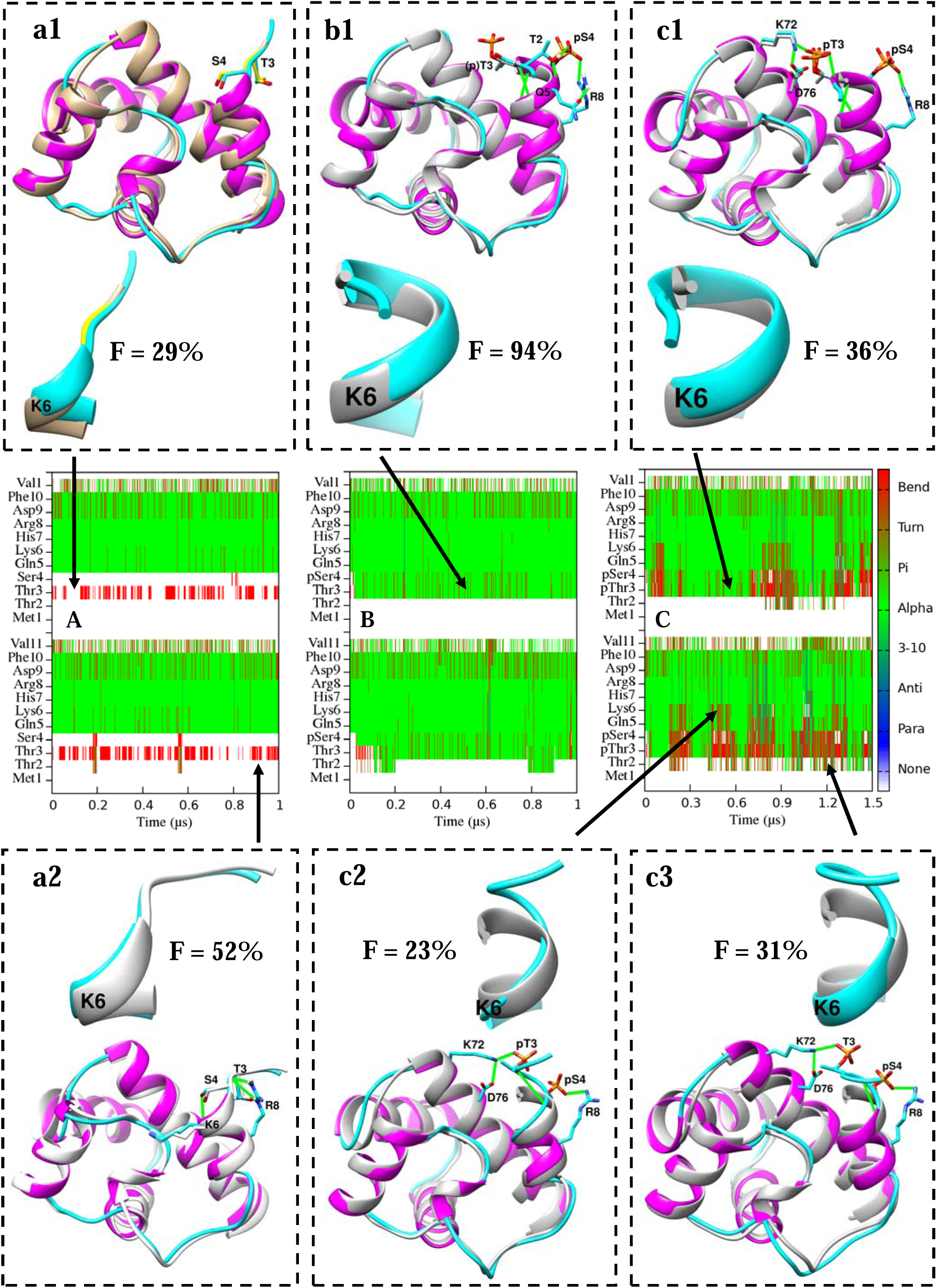
Secondary structure maps of the N-terminal residues 1-11 in two protomers of Banf1_Met1_ dimer and the corresponding Banf1_Met1_ representative conformations superimposed with reference structures. A, B, and C are the secondary structure maps of unphosphorylated Banf1_Met1_, mono-phosphorylated Banf1_Met1_, and di-phosphorylated Banf1_Met1_, respectively. a1 and a2 are the two representative conformations of unphosphorylated Banf1_Met1_. b1 is the representative conformation of mono-phosphorylated Banf1_Met1_. c1, c2, and c3 are the representative conformations of di-phosphorylated Banf1_Met1_. Zoom-in view of the N-terminal residues 1-7 of each representative structure is displayed in ribbon. Occupancies (F) of each representative conformation of Banf1 are also displayed accordingly. Conformations from MD are coloured by secondary structure, with helical structure in magenta and coil in cyan. The reference structure in a1 coloured in tan is adopted from chain A of 2BZF, with the phosphorylation sites of Thr3 and Ser4 highlighted in yellow. The reference structure in a2 coloured in white is adopted from chain A of 6UNT. Reference structures used for other conformations coloured in gray are chain A extracted from 7NDY. For clarity, sidechains of pSer4 and pThr3 in c2 and c3 are not displayed. Hydrogen bonds and salt bridges between charged residues are shown as green lines. Sidechains in reference structures are coloured in the same colour as their chain, whereas sidechains in MD representative structures are coloured in cyan. Oxygen, nitrogen, hydrogen and phosphorus atoms are coloured in red, blue, white and gold respectively.

**Figure S4.**
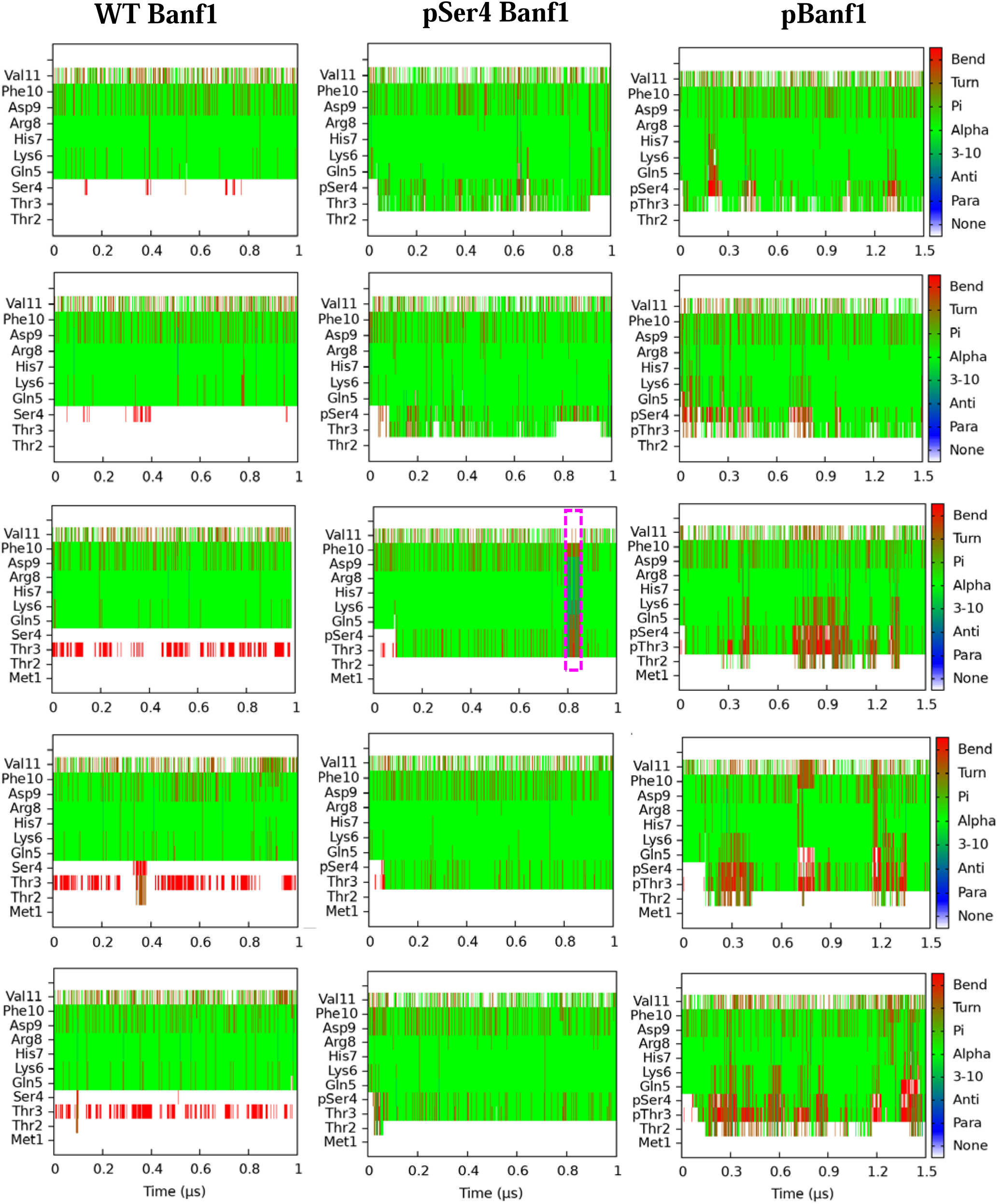
Secondary structure maps with respect to time of the N-terminal residues of Banf1, analysed from two simulation replicas of Banf1_Thr2_ monomer and three simulation replicas of Banf1_Met1_ monomer starting from different atomic velocities. The three secondary structure maps in the first two rows from left to right are for residues 2-11 of unphosphorylated, mono-phosphorylated and di-phosphorylated Banf1_Thr2_, respectively. The three secondary structure maps in the last three rows from left to right are for residues 1-11 of unphosphorylated, mono-phosphorylated and di-phosphorylated Banf1_Met1_, respectively. The representative secondary structure of pSer4 Banf1_Met1_ corresponds to the map in the magenta dashed line rectangular is displayed in Figure S5.

**Figure S5.**
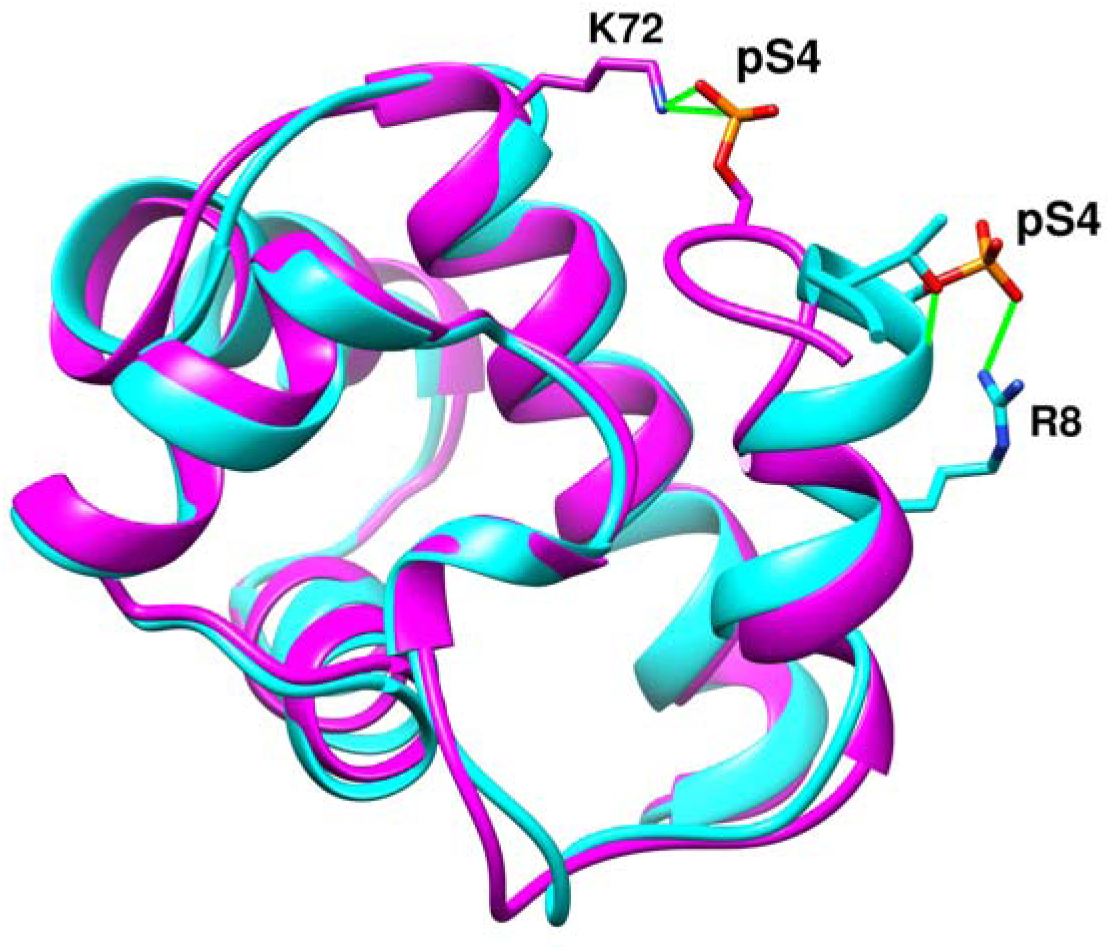
Overlay of the representative structure of pSer4 Banf1_Met1_ (magenta) corresponds to the secondary structure map in the magenta dashed line rectangular in Figure S4 and the representative structure of pSer4 Banf1_Met1_ (cyan) in Figure S3b1.

**Figure S6.**
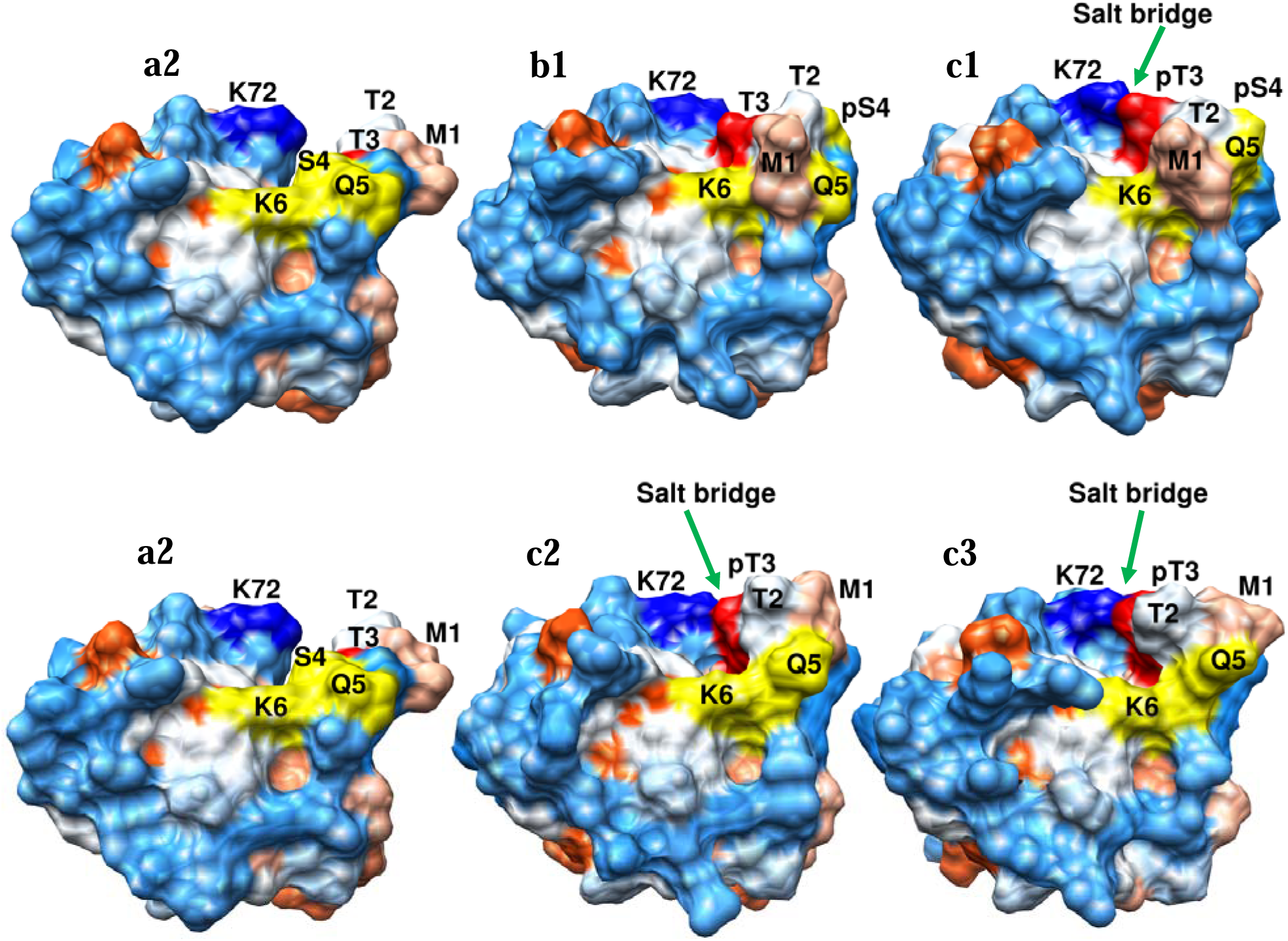
Hydrophobicity surface representation of the representative conformations of Banf1_Met1_ displayed in Figure S3, where blue indicates the surface is hydrophilic and red is for hydrophobic surface. a2 is for one of the representative conformations of WT Banf1_Met1_. b1 is for the representative conformation of pSer4 Banf1_Met1_. c1, c2, and c3 are for the three representative conformations of pBanf1_Met1_. Figure numbers for each representative conformation are consistent with those in Figure S3. N-terminal residues 4-6 involved in DNA binding are highlighted in yellow. Residue Met1, Thr2, (p)Thr3 and Lys72 are coloured in pink, white, red and blue respectively. The locations of salt bridges are directed by green arrows. In c2 and c3, surfaces of pSer4 are behind the surfaces of Met1, Thr2 and Gln5, and are not visible in this perspective.

**Figure S7.**
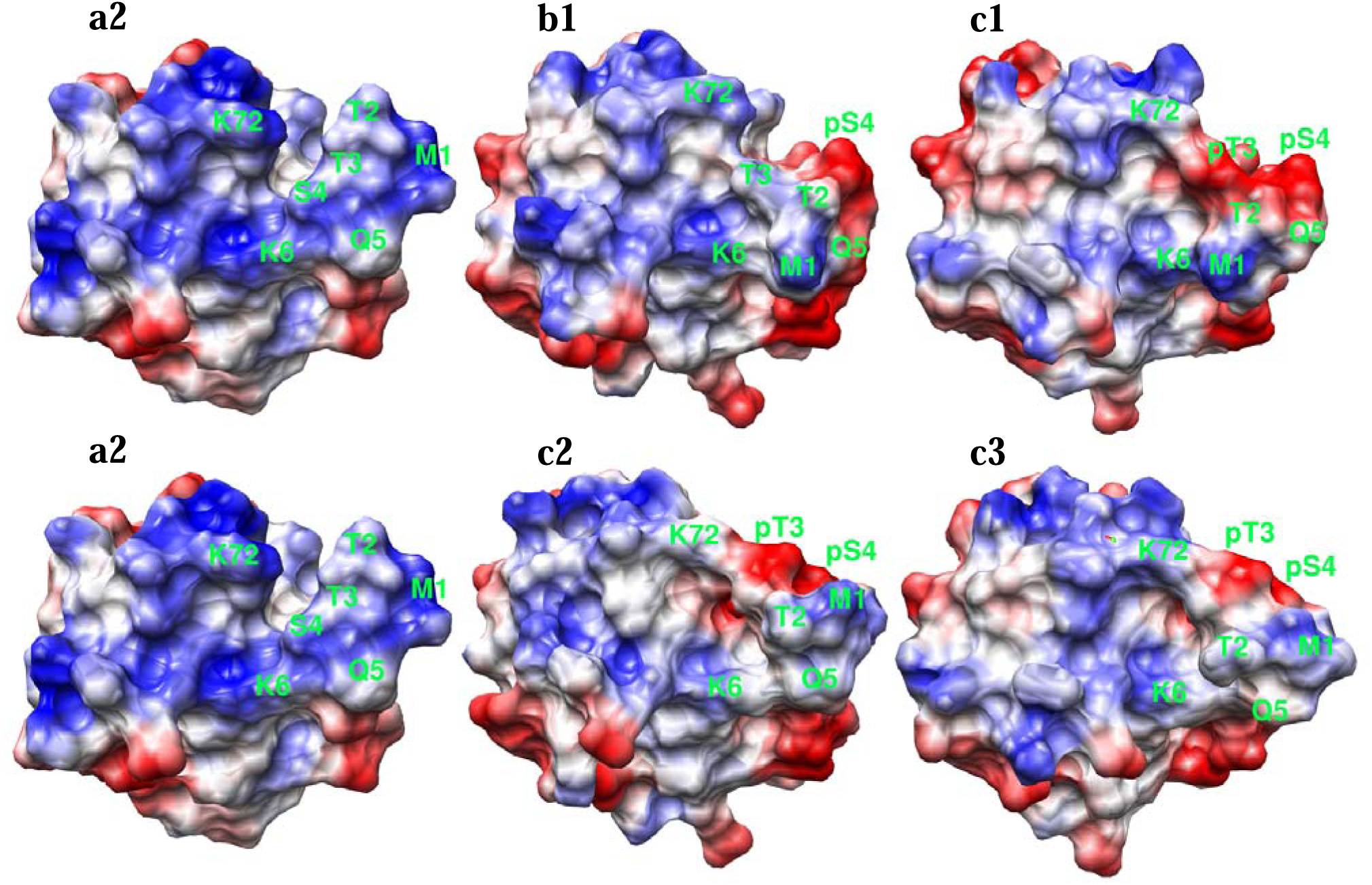
Electrostatic potential surface representation of the representative conformations of Banf1_Met1_ displayed in Figure S3 and S6, where blue, white and red correspond to electropositive, electroneutral, and electronegative surfaces, respectively. a2 is for one of the representative conformations of WT Banf1_Met1_. b1 is for the representative conformation of pSer4 Banf1_Met1_. c1, c2, and c3 are for the three representative conformations of pBanf1_Met1_. Figure numbers for each representative conformation are consistent with those in Figure S3 and S6.

**Figure S8.**
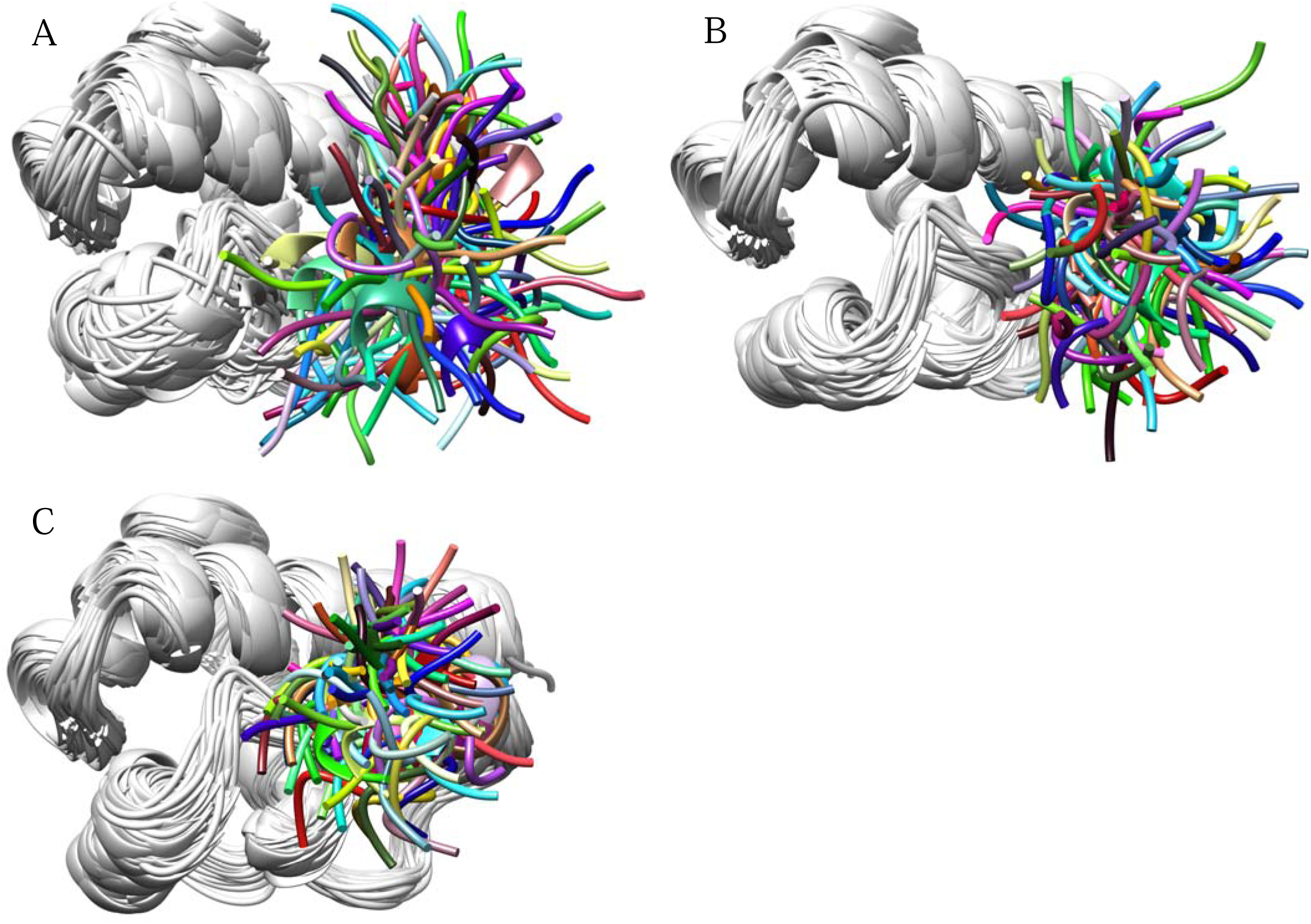
Top 100 representative conformation (in ribbon) of unphosphorylated (A), mono-phosphorylated (B) and di-phosphorylated (C) Banf1_Thr2_, where residues 8-89 are all coloured in gray and residues 2-7 are coloured differently in different conformations. The ensemble space occupied by Banf1 is restricted by mono-phosphorylation of Ser4. This restriction is more significant when pSer4 Banf1 is further phosphorylated at Thr3 forming pBanf1.

